# Identification of uranyl-binding proteins in *Arabidopsis thaliana* cells exposed to uranium: Insights from a metalloproteomic analysis and characterization of Glycine-Rich RNA-binding Protein 7 (GRP7)

**DOI:** 10.1101/2025.04.28.650916

**Authors:** Benoit H. Revel, Adrien Favier, Jacqueline Martin-Laffon, Alicia Vallet, Jonathan Przybyla-Toscano, Sabine Brugière, Yohann Couté, Hélène Diemer, Sarah Cianférani, Thierry Rabilloud, Jacques Bourguignon, Bernhard Brutscher, Stéphane Ravanel, Claude Alban

## Abstract

Uranium (U) is a naturally occurring radionuclide, chemotoxic for living organisms. To identify proteins that could be cellular targets for U in plants, we used metalloproteomic approaches combining column chromatographic fractionation analyses, protein identification by high-resolution mass spectrometry shotgun proteomics, and metal quantification by induced coupled plasma mass spectrometry. We identified 57 candidate proteins for uranyl (U(VI)) binding in cultured *Arabidopsis thaliana* cells. One of these proteins, the Glycine-Rich RNA-binding Protein 7 (GRP7) is a RNA-binding protein involved in various developmental processes and responses to biotic and abiotic stress. Recombinant GRP7 was purified from overproducing bacteria and subjected to further biochemical characterization. First, we showed that GRP7 binds U(VI) with a 1:2 (protein:metal) stoichiometry *in vitro*. Next, we analyzed its structural properties by solution-state nuclear magnetic resonance spectroscopy. This allowed us to gain insight into the molecular dynamics of the protein-metal interaction and to identify residues involved in U(VI)-binding in the two sites. Finally, we observed that U(VI) binding interferes with nucleic acid binding to the protein RNA-binding domain, suggesting that U(VI) binding to GRP7 contributes to U toxicity in plants.

**HIGHLIGHTS:**

- 57 candidate uranyl-binding proteins were identified from Arabidopsis cells exposed to uranyl, of which the RNA-binding protein GRP7.
- *In vitro*, recombinant GRP7 binds 2 uranyl ions within the RNA recognition motif domain.
- Amino acid residues involved in uranyl binding in both binding sites were identified.
- Competition between specific oligonucleotide and uranyl for binding suggests the implication of GRP7 in uranium toxicity, as a cellular target.

## 1- Introduction

Uranium (U) is a non-essential trace metal element that is ubiquitous in the Earth crust and seawater. The radionuclide is primarily redistributed in the environment by anthropogenic activities including U mining and milling industries, civil and military nuclear activities, and extensive enrichment of agricultural soils with phosphate fertilizers. Its accumulation in soil, water and air can lead to potential risks to ecosystems, agrosystems, and ultimately human health, as the radionuclide has both chemical and radiological toxic effects [1].

The soluble oxidized form of U, the uranyl cation (UO_2_^2+^, with U in the +VI oxidation state, thereafter referred to as U(VI)), is known to induce oxidative stress, to affect photosynthesis, to disrupt mineral homeostasis and to inhibit plant growth and development, including root architecture and elongation [2–11]. U(VI) has also been shown to alter the expression of genes involved in hormone synthesis and signaling, plasmodesmata function, and cell wall metabolism [4, 12–14]. More recently, a proteomic study of root membrane and cell wall revealed the importance of vesicular trafficking and water balance in the response of plants to U(VI) exposure [15]. Despite these studies, our knowledge of the molecular mechanisms responsible for U toxicity in plants is still fragmentary. Toxic effects of U on organisms result from direct interactions with biological molecules. Since U(VI) is able to bind strongly to biomolecules *via* carboxylate, phosphate or sulphate moieties, proteins are expected to be primary targets of U [16]. These interactions also determine the molecular mechanisms of U(VI) uptake, transport, and storage. Hence, we have recently shown that U(VI) is taken up by roots of *Arabidopsis thaliana* through calcium channels [17]. Calcium channels, but also iron permease, are involved in U(VI) uptake in yeast [18]. Another pathway involving endocytic uptake may also be important for U(VI) transport into plant cells [19]. However, once inside cells, the fate of U(VI) and the nature of its protein targets are still poorly understood. We also do not know whether plants are able to synthesize peptides/proteins that could be involved in U detoxification. Such peptides/proteins, *e.g.* phytochelatins and metallothioneins, are present in the cell for its protection against an excess of metals such as cadmium [20].

In recent years, a growing number of studies have focused on the identification and characterization of protein-U(VI) interactions *in vivo* and *in vitro* in various organisms, allowing the identification of potential U(VI) targets *in vivo* (reviewed in [21] and [22]; [23]). In order to identify U(VI)-binding proteins in mammals, Vidaud and colleagues have developed an immobilized uranyl affinity chromatography (IMAC) method, based on the cation-exchange properties of aminophosphonate groups for U(VI) binding [24]. This IMAC technique was a powerful tool for the *in vitro* capture of U(VI)-binding proteins (UraBPs) from human serum [24] and from human kidney-2 cell extracts [25]. Recently, we have made a major breakthrough in U toxicity in a photosynthetic organism by identifying the first set of UraBPs from *A. thaliana* root and shoot extracts using the above-mentioned modified IMAC technique [26]. However, although the plasma membrane-associated cation-binding protein PCaP1, identified in this study as an *in vitro* UraBP, was shown to influence U(VI) translocation from roots to shoots *in planta*, its role in this process remains to be fully understood. Moreover, the identification of proteins with *in vitro* U(VI) affinity using such a chromatographic method does not inherently indicate their involvement in U toxicity or detoxification processes *in vivo*.

In a previous work, we developed an ionomic, metalloproteomic, and biochemical toolbox to analyse the consequences of U(VI) ion stress on the proteome of *A. thaliana* cultured cells [27]. We showed that high-resolution fractionation of *A. thaliana* cell soluble proteins by anion-exchange chromatography (AEC) coupled to MS-based proteomics is a very efficient method, invaluable for reducing sample complexity and improving proteome coverage through enrichment of low-abundant proteins. Uranium was detected in several chromatographic fractions, indicating that U(VI)-protein interactions were at least partially preserved during fractionation. This finding also provided the first evidence that several *A. thaliana* proteins can bind U(VI) *in vivo*.

Here, we aim to identify *A. thaliana* proteins with *in vivo* affinity for the U(VI) cation. A key challenge was to separate U(VI)-bound proteins using chromatographic techniques that preserve metal-protein interactions for accurate identification. To address this, we first refined the conditions for exposing cell cultures to U(VI), ensuring the stability of protein-metal complexes. We then implemented two complementary fractionation strategies, combining sequential column chromatography, ICP-MS metal profiling and high-resolution MS-based shotgun proteomics. This enabled us to identify 57 UraBP candidates from *A. thaliana* cultured cells exposed to U(VI) contamination. One of these proteins, the glycine-rich RNA-binding protein 7 (GRP7), which was one of most highly enriched of the overall process was further characterized for its ability to interact with U(VI) using a combination of biochemical and structural analyses. Taken together, our results showed that U(VI) binds to the recombinant GRP7 core domain at two distinct sites through intramolecular interactions. We identified important amino acids for U(VI) binding within both sites. Finally, we investigated the relationship between U(VI)-binding and nucleotides binding to GRP7 protein.

## 2- Materials and Methods

### 2.1. Arabidopsis thaliana cell culture growth conditions

*Arabidopsis thaliana* (ecotype Columbia) cell cultures were grown in Murashige and Skoog liquid medium supplemented with 1.5% sucrose (w/v), at 22°C, under continuous light (40 µmol photons m^-2^s^-1^) [27]. Cells were subcultured every 4 days for 3 consecutive cycles before exposure to U(VI). To this end, exponentially growing cells were harvested by centrifugation, washed once and resuspended in medium with low (30 µM) or no phosphate at all, instead of 1.25 mM in regular medium. Cells were then challenged with 50 μM uranyl nitrate for 24 h. After exposure, cells were harvested, washed once with 10 mM sodium carbonate solution followed by two washes with distilled water, dried by vacuum filtration, and stored at -80°C prior to use [27].

### 2.2. Extraction of soluble proteins from cell cultures

*A. thaliana* cultured cells exposed with U(VI) were ground in liquid nitrogen using a mortar and pestle. The powdered samples were suspended in 10 mM TrisHCl buffer pH 7.5, 1 mM dithiothreitol (DTT) and a protease inhibitor cocktail (Roche Applied Science). Suspensions were then centrifuged at 16,000 x g for 20 min at 4°C. Supernatants, comprising soluble proteins, were recovered, ultra-centrifuged at 105,000g for 15 minutes at 4°C, and desalted by successive concentrations/dilutions with 10mM TrisHCl pH 7.5, 1 mM DTT buffer (Strategy 1) or 20 mM K_2_HPO_4_/KH_2_PO_4_ pH 7.5, 1 mM DTT buffer (Strategy 2), using Amicon® Ultra-15, 3 kD ultrafiltration units (Millipore).

### 2.3. Fractionation of soluble cell proteins by column chromatography

All chromatographic steps were carried out at 4°C using Fast Protein Liquid Chromatography using an Äkta purifier (Cytiva).

### Strategy 1

#### First step – Anion exchange chromatography (AEC) fractionation

Aliquots of 400 mg soluble proteins from *A. thaliana* cells exposed to U(VI) were loaded onto a 1.6 x 12 cm Q-Sepharose High Performance column (Cytiva) equilibrated with 10 mM Tris-HCl pH 7.5, 1 mM DTT (buffer A) at a flow rate of 0.5 ml/min. Proteins were then eluted using a linear gradient of NaCl to 0.6 M, in buffer A over 14 column volumes. Fractions of 5 ml were collected.

#### Second step – Size exclusion chromatography (SEC) fractionation

Protein fractions from the AEC fractionation, co-eluting with U, were concentrated, loaded onto a Superdex 200 HiLoad® 16/60 column (Cytiva) equilibrated with 10 mM Tris-HCl buffer pH 8, 10% (v/v) glycerol, 0.15 M NaCl at a flow rate of 1 ml/min, and then eluted with the same buffer. Fractions of 1.5 ml were collected.

### Strategy 2

#### First step - Pseudo affinity chromatography fractionation

Aliquots of 100 mg of soluble proteins from *A. thaliana* cells exposed to U(VI) were fractionated by chromatography onto a 5 ml HTP grade II hydroxyapatite column (Bio-Rad) equilibrated with 20 mM K_2_HPO_4_/KH_2_PO_4_ pH 7.5 phosphate buffer, 1 mM DTT. Elution was performed with a linear gradient of K_2_HPO_4_/KH_2_PO_4_ pH 7.5 from 20 mM to 1 M, over 20 column volumes. Fractions of 5 ml were collected.

#### Second step - Size exclusion chromatography (SEC) fractionation

The HTP fractions containing the major U peak, eluted by the phosphate gradient, were pooled and concentrated using Amicon Ultra-4, 3 kDa filtration units (Millipore). The sample was then chromatographed onto a Superdex 200 HiLoad® 16/60 column (Cytiva) equilibrated with 10 mM Tris-HCl buffer pH 8, 10% (v/v) glycerol, 0.15 M NaCl at a flow rate of 1 ml/min, and eluted with the same buffer. Fractions of 1.5 ml were collected.

#### Third step – Anion exchange chromatography (AEC) fractionation

The fractions of the five U peaks eluted from the SEC step were pooled and diluted in 10 mM Tris-HCl pH 8 buffer to lower salt concentration. The samples were then chromatographed onto a 1 ml Q-Sepharose High Performance column (Cytiva), using a linear gradient of NaCl up to 0.6 M, over 14 column volumes. Fractions of 400 µl were collected.

### 2.4. Protein determination and SDS-PAGE analyses

Protein were quantified by the Bradford method using Bio-Rad protein assay reagent, with bovine serum albumin as a standard [28]. Aliquots of protein fractions were separated on 12% or 15% (w/v) polyacrylamide gels under denaturing conditions (SDS-PAGE) and stained with Coomassie Blue using InstantBlue^TM^ protein stain (Expedeon) or silver nitrate using the SilverQuest staining kit (Invitrogen), according to the supplier’s recommendations.

### 2.5. ICP-MS analyses

Protein fractions from chromatographic columns were mineralized in a 10% (v/v) HNO_3_ solution (Suprapur; Merck) by heating for 2 h at 60°C to ensure denaturation and metal release [26, 27]. Denatured proteins were removed by centrifugation and the supernatants used for ICP-MS analysis. The mineralised samples were diluted in a 0.5% (v/v) HNO_3_ solution and then analyzed using an iCAP RQ quadrupole mass spectrometer (Thermo Fisher Scientific GmbH, Germany) equipped with a MicroMist U-Series concentric glass nebuliser, a 3°C cooler, a Qnova quartz torch, two nickel cones with a high-sensitivity insert coupled to an ASX-560 autosampler (Teledyne CETAC Technologies, USA). Uranium was quantified using standard mode and standard curves, as well as an internal standard solution containing ^103^Rh and ^172^Yb. Data integration was performed using Qtegra software (Thermo Fisher Scientific GmbH, Germany).

### 2.6. Mass spectrometry-based proteomic analyses (UraBPs identification; Strategy 1)

The 2D gel-based proteomic experiments were essentially carried out as previously described [29]. For the isoelectric focusing (IEF) dimension, home-made 160 mm long 4-8 linear pH gradient gels were used. Four mm-wide strips were cut, and rehydrated overnight with the sample, diluted in a final volume of 0.6 ml of rehydration solution (7 M urea, 2 M thiourea, 4% CHAPS, 0.4% carrier ampholytes (Pharmalytes 3-10) and 100 mM dithiodiethanol [30]). The strips were then placed in a Multiphor plate (GE Healthcare), and IEF was carried out with the following electrical parameters: 100V for 1 h, then 300 V for 3 h, then 1000 V for 1 h, then 3400 V up to 60-70 kVh. After IEF, the gels were equilibrated for 20 min in 125mM Tris, 100mM HCl, 2.5% SDS, 30% glycerol and 6 M urea. They were then transferred on top of the SDS gels and sealed in place with 1% agarose dissolved in Tris 125mM, HCl 100mM, SDS 0.4% and 0.005% (w/v) bromophenol blue.

For the second dimension, 10% acrylamide gels (160×200×1.5 mm) were used. The Tris taurine buffer system [31] was used and operated at a ionic strength of 0.1 and a pH of 7.9. The final gel composition is thus Tris 180mM, HCl 100 mM, acrylamide 10% (w/v), bisacrylamide 0.27%. The upper electrode buffer is Tris 50 mM, taurine 200 mM, SDS 0.1%. The lower electrode buffer is Tris 50 mM, glycine 200 mM, SDS 0.1%. The gels were run at 25 V for 1h, then 12.5 W per gel, until the dye front has reached the bottom of the gel. Detection was carried out by a tetrathionate silver staining [32].

The gels were scanned after silver staining on a flatbed scanner (Epson perfection V750), using a 16 bits grayscale image acquisition. The gel images were then analyzed using the Delta 2D software (v 3.6). In order to select potential UraBPs, a protein correlation strategy was used: several fractions corresponding to the various U-containing peaks were analyzed by 2D electrophoresis, taking care of including the fraction with the highest U content and other fractions at different heights of the U-content scale. Spots showing the highest level in the fraction with the highest U content were selected as potential U-binding proteins.

Spots excision, destaining and in gel digestion were performed as described in [33]. The peptide digests were then analysed by nanoLC-MS/MS using a nanoACQUITY Ultra-Performance-LC (Waters Corporation, Milford, USA) coupled to the TripleTOF 5600 (Sciex, Ontario, Canada) operated as described in [34].

For protein identification, the MS/MS data were interpreted using a local Mascot server with MASCOT 2.6.2 algorithm (Matrix Science, London, UK) against UniProtKB/SwissProt database (version 2019_09 561,176 sequences and version 2019_10 561,356 sequences). Spectra were searched with a mass tolerance of 15 ppm for MS and 0.07 Da for MS/MS data. Trypsin was selected as the enzyme allowing a maximum of one missed cleavage. Carbamidomethylation of cysteine residues and oxidation of methionine residues were specified as variable modifications. Protein identifications were validated with at least two peptides with Mascot ion score above 30.

### 2.7. Mass spectrometry-based proteomic analyses (UraBPs identification; Strategy 2)

Proteins were solubilized in Laemmli buffer before being stacked in the top of a 4-12% NuPAGE gel (Invitrogen). After staining with R-250 Coomassie Blue (Biorad), proteins were digested in-gel using trypsin (modified, sequencing purity, Promega), as previously described [35]. The resulting peptides were analyzed by online nanoliquid chromatography coupled to MS/MS (Ultimate 3000 RSLCnano and Q-Exactive Plus) using a 80 min gradient. For this purpose, peptides were sampled on a precolumn (300 μm x 5 mm PepMap C18, Thermo Scientific) and separated in a 75 μm x 250 mm C18 column (Reprosil-Pur 120 C18-AQ, 1.9 μm, Dr. Maisch). The MS and MS/MS data were acquired by Xcalibur v2.8 (Thermo Fisher Scientific).

Peptides and proteins were identified by Mascot (version 2.8.0, Matrix Science) through concomitant searches against the Uniprot database (Arabidopsis thaliana taxonomy, January 2024 version), and a homemade database containing the sequences of classical contaminant proteins found in proteomic analyses (human keratins, trypsin, etc.). Trypsin/P was chosen as the enzyme and two missed cleavages were allowed. Precursor and fragment mass error tolerances were set at respectively at 10 and 20 ppm. Peptide modifications allowed during the search were: Carbamidomethyl (C, fixed), Acetyl (Protein N-term, variable) and Oxidation (M, variable). The Proline software ([36], version 2.2) was used for the compilation, grouping, and filtering of the results (conservation of rank 1 peptides, peptide length ≥ 6 amino acids, false discovery rate of peptide-spectrum-match identifications < 1% [37], and minimum of one specific peptide per identified protein group).

Proteins from the contaminant database were discarded from the final list of identified proteins. Proteins identified with at least 2 peptides in the two samples analyzed and proteins with molecular weight inferior to 20 kDa were further considered.

### 2.8. Cloning of GRP7 cDNA forms for recombinant proteins production and site-directed mutagenesis

Sequences of primers used in this study are listed in Supplementary Table S1. The full-length GRP7 cDNA coding sequence (At2g21660) was amplified by RT-PCR using total RNA isolated from 3-week-old *A. thaliana* leaves using the RNeasy plant mini extraction kit (Qiagen) and reverse transcribed using the ThermoScript RT-PCR system (Invitrogen). The cDNA fragment was amplified by PCR using primers introducing *Nco*I and *Sal*I restriction sites upstream of the initiation codon and downstream of the stop codon, respectively (Supplementary Table S1). The PCR products were sequenced (Eurofins) and ligated into pET28b+ plasmid (Novagen) between *Nco*I and *Sal*I sites to obtain the recombinant pET28-GRP7 plasmid. cDNA encoding the C-terminal truncated GRP7 form, GRP7Δ, was obtained by PCR amplification using pET28-GRP7 plasmid as a matrix and specific primers (Supplementary Table S1). The PCR product was ligated into pET28b+ plasmid to obtain the pET28-GRP7Δ recombinant plasmid. The resulting pET28-GRP7 constructs were amplified in *Escherichia coli* DH5α cells and then introduced into the *E. coli* overexpression host Rosetta 2 (DE3) (Stratagene).

Mutations into GRP72 were introduced by a PCR-based strategy according to the instructions of the QuickChange II site directed mutagenesis kit (Stratagene). For the generation of triple mutants, double mutant constructs were used as templates (for primers, see Supplementary Table S1).

### 2.9. Production and purification of the recombinant GRP7 forms

Bacterial cells transformed with GRP7 plasmid constructs were cultured at 37°C in Lysogeny Broth (LB) medium supplemented with the appropriate antibiotics until A_600_ reached 0.6. Isopropylthio-β-D-galactoside was then added to a final concentration of 0.4 mM and incubations were continued for 15 h at 28°C. Cell pellets from 2-L cultures were resuspended in 50 ml of extraction buffer, which contained 20 mM Tris-HCl, pH 8, 10% (w/v) glycerol, 1 mM DTT and a cocktail of complete protease inhibitors (Roche Applied Science), and were then disrupted by sonication using a Vibra-Cell disruptor (Branson Ultrasonics). The removal of cell debris was achieved through centrifugation at 40,000 x *g* for 30 min. The soluble proteins present in the supernatants were then subjected to ammonium sulphate precipitation at 4°C with crystalline ammonium sulphate ranging from 40 to 60% saturation, for GRP7 containing protein extracts, or 60 to 90% saturation, for GRP72 containing protein extracts. Resulting precipitates were collected by centrifugation (40,000 x *g*, 15 min, 4°C), resuspended in 20 mM Tris-HCl, pH 7.5 buffer, supplemented with protease inhibitors, and dialyzed overnight at 4°C against 4 L of the same buffer. Protein samples were applied onto a 1.6 x 10 cm Q-Sepharose High Performance column (Cytiva) equilibrated with 20 mM Tris-HCl, pH 7.5 buffer. After extensive wash of the column with two volumes of buffer, proteins were eluted using a 11-column volume linear gradient from 0 to 300 mM NaCl in this buffer, at a flow rate of 0.5 ml/min. GRP7-containing fractions were pooled and concentrated using Amicon® Ultra-15, 3kDa filtration units (Millipore), before being applied onto a Hiload® Superdex 75 16/60 column (Cytiva), equilibrated with 10 mM Tris-HCl, pH 7.5, 150 mM NaCl buffer. Elution was conducted in the same buffer at a flow rate of 1 ml/min. Purified recombinant proteins were concentrated, aliquoted and stored at -80°C, until use. Protein purity and integrity were monitored by SDS-PAGE. Purified protein concentration was determined by recording UV absorption spectra, using a NanoDrop 2000 spectrophotometer (Thermo Fischer Scientific) (mass extinction coefficients at 280 nm were E_1%_ = 15.33 for GRP7; 8.68 for GRP72; 8.77 for GRP72U1_mut_ and GRP72U2_mut_; and 8.86 for GRP72U1-U2_mut_, as calculated from molar extinction coefficients c_280_ = 25900 M^-1^ cm^-1^ (GRP7) and 8480 M^-1^ cm^-1^ (GRP72 and mutants); ProtParam ExPASy). Production and purification of recombinant GRP7 and GRP72 proteins uniformly labelled with ^15^N and ^13^C (*U*-^15^N,^13^C-GRP7) were performed similarly as for unlabeled protein, except that growth was conducted in minimal M9 medium, containing antibiotics and supplemented with^15^N-ammonium chloride (^15^N-NH_4_Cl (^15^N, 99%)) and ^13^C-glucose (*U*-^13^C6, 99%) (Cambridge Isotope Laboratories, Inc.), after progressive medium acclimatization from pre-cultures conducted in LB medium.

Presence of recombinant GRP7 in eluted fractions during the purification process was assessed by western-blotting, using a custom-made rabbit polyclonal GRP7 antibody raised against purified recombinant 6His-tagged protein (1:25000 dilution) (Covalab).

### 2.10. Uranium quantification in U(VI)-protein complexes by Size Exclusion Chromatography (SEC) and Arsenazo III assay

U(VI)-protein complexes were prepared by incubating 15 μM purified GRP7 solutions for 15-30 min at 25°C, in 20 mM Tris-HCl pH 7.5 buffer, containing 150 mM NaCl, 150 µM iminodiacetic acid and up to 200 μM uranyle nitrate, as indicated. Iminodiacetic acid was present in incubation buffer to prevent U(VI) precipitation as hydro-U(VI) hydrolysates [38]. Formed complexes were separated from unbound metal by SEC, through centrifugation for 2 min at 724 x *g* of the samples (120 μL), on MicroSpin™ G-25 columns (Cytiva), equilibrated with the binding buffer [26]. Controls without protein or without U(VI) were run in parallel for background correction. Binding assays were performed at least in triplicate. Protein in the eluates was quantified by recording A_280_. Uranium in protein complexes was determined in 100 µL aliquots by the Arsenazo III colorimetric assay [39], on ELISA plates, using a microplate reader (Infinite M1000 PRO, TECAN) for detection. The reliability of the Arsenazo III method for U quantification was confirmed by comparing to an ICP-MS assay, using a standard range of 0-30 µM uranyl nitrate.

### 2.11. Statistical analyses

Statistical analysis of the data in U(VI) binding experiments was performed using Dunnett-test in Kaleidagraph 5.0 (Synergy Software, PA, USA). A *P*-value<0.05 was considered statistically significant.

### 2.12. Solution-state Nuclear Magnetic Resonance (NMR) studies

Uniformly ^13^C/^15^N labeled GRP7 (or GRP72) was dissolved in the NMR buffer consisting in Tris-HCl 20 mM (pH 7.5), NaCl 0.3 M, and placed in either 3 mm or 5 mm NMR sample tubes. The final sample concentration ranged between 100 and 200 µM, and the sample temperature was set to 300 K for the NMR experiments.

NMR experiments were performed on Bruker Avance IIIHD spectrometers operating at ^1^H Larmor frequencies of 700, 850 and 950 MHz. All spectrometers were equipped with cryogenically cooled triple-resonance probes (HCN TCI 5mm) and pulsed z-field gradients. 2D ^1^H-^15^N and ^1^H-^13^C correlation spectra were recorded with BEST-TROSY [40] and SE-HSQC pulse sequences, respectively.

NMR assignments were obtained from a set of 3D BEST-TROSY-type correlation experiments [41]: HNCO, HNCACO, HNCA, HNCOCA, HNCACB, and HNCOCACB.

Translational diffusion constants of the proteins in solution were measured by 1D methyl ^1^H DOSY experiments [42] focusing either on the methyl or amide spectral region. A series of 1D spectra was recorded with varying gradient strength and a total acquisition time of about 15 min.

All NMR experiments used in this study are implemented in the NMRlib pulse sequence library [43] that can be freely downloaded from the IBS website (http://www.ibs.fr/research/scientific-output/software/pulse-sequence-tools). The experiments were processed and analyzed using Bruker Topspin 3.5 and CCPNMR V3 software.

## 3- Results

### 3.1. Optimisation of U(VI) accumulation in A. thaliana cells and strategies used to identify in cellulo U(VI)-binding proteins (UraBPs)

The metalloproteomic and biochemical strategy that we developed to analyze the consequences of U(VI) stress on the proteome of *A. thaliana* cell cultures [27] was used as a starting point to initiate the purification of *in cellulo A. thaliana* U(VI)-binding proteins (UraBPs). In this work, *A. thaliana* cells growing exponentially in standard MS medium were transferred to a cell density of 100 g fresh weight/L in low phosphate (30 µM instead of 1.5 mM) MS medium and were challenged with 50 µM uranyl nitrate for 24 h. Under these conditions, the proportion of soluble U measured in cells was low (1.4 ± 0.8 % from 71 ± 20 µg U biosorbed/ g fresh weight) and the majority of U was found as an insoluble form adsorbed on cell walls (60-70%) and biological membranes (20-30%) [27]. In order to maximize U(VI) uptake and accumulation into cells and to favor intracellular U(VI) binding to proteins, we made adjustments to the original experimental protocol. Phosphate being known to limit U(VI) absorption [5, 44], it was completely omitted prior to U(VI) exposure. Also, the cell density was lowered to 20-30 g fresh weight/L increasing cell exposure to U(VI). Under these conditions, the exposure of cells to 50 µM uranyl nitrate did not affect cell growth, and total U biosorbed was 156.8 ± 14.1 µg/g of fresh cell weight (Supplementary Figure S1). Cells were harvested after 24 h of treatment and extensively washed with Na_2_CO_3_ and distilled water to remove U(VI) loosely adsorbed to the cell surface. Cells were then lysed and ultracentrifuged to separate the supernatant, corresponding to the soluble protein fraction, from the insoluble fraction containing cell wall and membranes. Uranium quantification by ICP-MS showed that 83 ± 3 % of U was present in the insoluble fraction confirming that most of the radionuclide was associated with the cell wall, membranes or precipitates eliminated by ultracentrifugation [27]. The soluble protein fraction that contained 17 ± 3 % of total U(VI) biosorbed was then desalted by successive ultrafiltration and dilution (with a 3 kDa cut-off membrane) in order to retain only U(VI) associated with proteins. The final soluble protein extract that contained 169 ± 13 ng U/ mg of protein (vs 25 ± 12 ng U/ mg protein under low phosphate conditions [27]) was used to purify and identify *A. thaliana* UraBPs using MS-based proteomic approaches. To achieve this goal, two distinct biochemical strategies were developed, both relying on the successive fractionation of the soluble protein extract using different chromatographic columns (Figure 1). The first strategy aimed at the comprehensive identification of UraBPs, while the second focused on isolating proteins with the highest affinity for U(VI).

**Figure 1.**
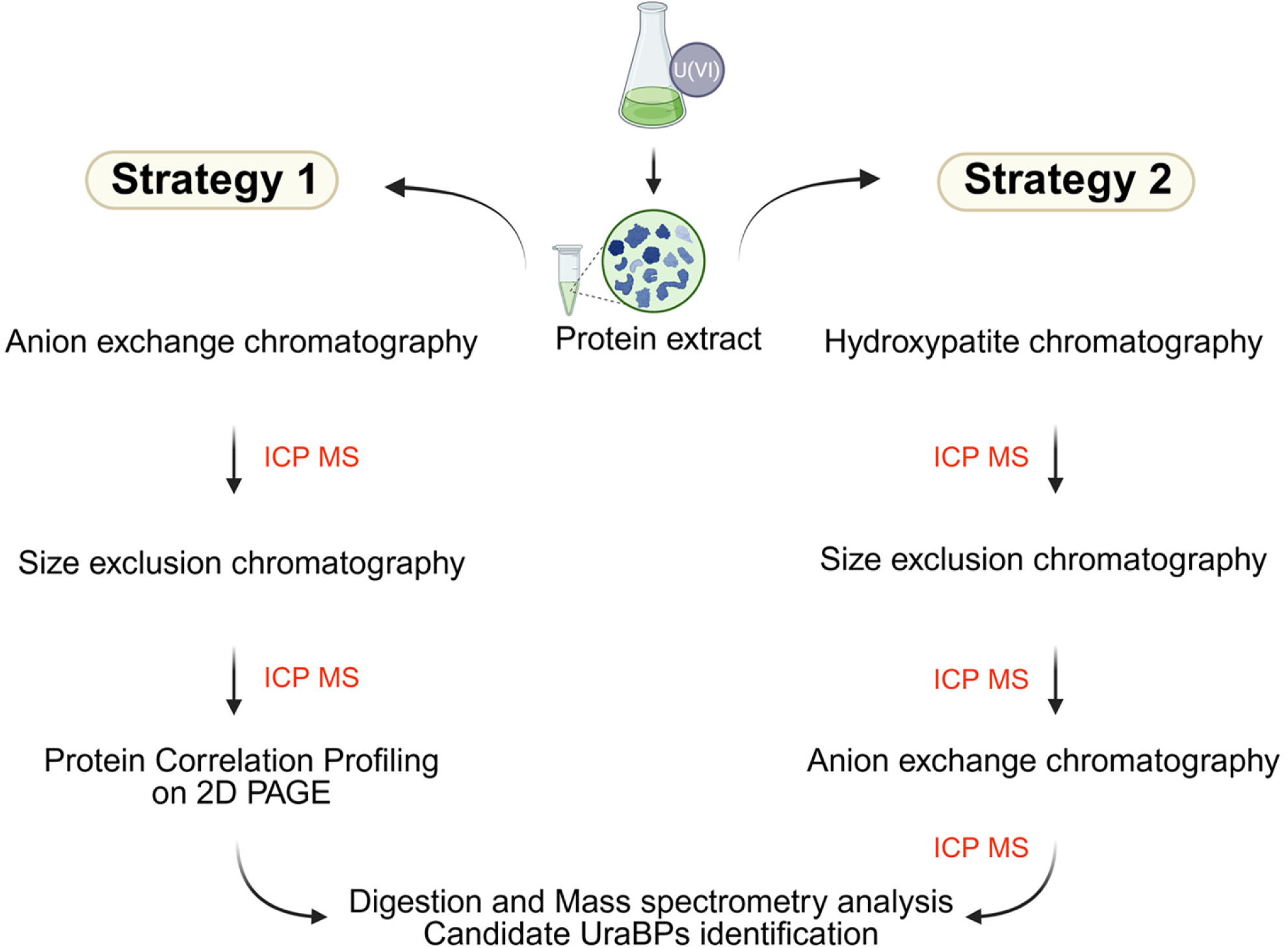
Strategies used for the identification of *in cellulo* U(VI)-binding proteins (UraBPs). Two sequential chromatographic strategies were used to identify candidate UraBPs from soluble proteins of *A. thaliana* cells exposed to uranyl nitrate. Uranium was quantified at each chromatographic step using ICP-MS. Strategy 1 prioritized the preservation of ionic bonds for a comprehensive UraBP analysis. It involved three steps: (1) High resolution Q-Sepharose anion exchange chromatography, (2) size exclusion chromatography on a Superdex 200 column, and (3) 2D-gel differential analysis of fractions collected from the middle and top of the eluted U peaks. Strategy 2 focused on isolating proteins with the highest affinity for U. It began with hydroxyapatite chromatography to remove low-affinity proteins, followed by Superdex 200 size exclusion chromatography and a final high-resolution step on a Q-Sepharose column. Candidate UraBPs were identified by nLC-MS/MS.

### 3.2. UraBP identification by Strategy 1: Towards a comprehensive identification of the U(VI)-binding proteins in cellulo

In strategy 1, three grams of soluble proteins extracted from 200 g of *A. thaliana* cells were separated by anion exchange chromatography (AEC, Q-Sepharose HP column) using a continuous NaCl gradient from 0 to 0.6 M (Figure 1). The presence of U in the fractions eluted from the AEC column was measured by ICP-MS. We identified five distinct peaks containing U, named peak 1 to 5 (Figure 2A). These peaks were eluted in a highly reproducible manner as they were observed at each chromatography iteration, with the exception of peak 5 whose intensity varied from one experiment to another (not shown). This result showed that there are different populations of UraBPs in *A. thaliana* cells challenged with uranyl nitrate. In these five peaks, the protein composition was very different and still very complex (Figure 2B), suggesting that each of these peaks may contain several UraBPs and possibly associated proteins. The composition of peak 5 was different from the other four peaks because, in addition to containing few proteins (Figure 2B), most of the U present in this peak was in the free form or bound to small molecules (< 3 kDa). In fact, when peak 5 was concentrated on a centrifugal filter unit (3 kDa cut off), 80 % of U was not retain by the filter, whereas in peaks 1 to 4, this proportion represented only 5 to 10 %. Accordingly, peak 5 was not analyzed further in this study. For further refinement, a Superdex 200 size-exclusion chromatography (SEC) column was employed due to its effectiveness in maintaining U(VI)-protein complexes [45–48]. This step revealed that individual AEC-derived U peaks contained multiple UraBPs. Indeed, Peaks 1 and 2 from the AEC further separated into 3 and 4 distinct U peaks, respectively (named 1.I to 1.III and 2.I to 2.IV), whereas Peaks 3 and 4 from the AEC generated only one main U peak (3.I and 4.I, respectively) eluted from the Superdex 200 column exclusion volume (Supplementary Figure S3). However, the high complexity of polypeptide profiles on SDS-PAGE, combined with the significant decline in the U-to-protein ratio observed after these two chromatographic steps complicated further separation using this approach. To address this limitation, we developed an alternative strategy integrating 2D gel electrophoresis and protein correlation profiling. We compared the 2D protein maps of the peak fractions (highest U content) with those of the flanking fractions eluted from the chromatographic column before and after the peak maximum. Correlating protein abundance with U enrichment enabled the identification of potential UraBP candidate spot proteins. The candidate UraBPs were then identified by nLC-MS/MS. This approach was tested on two SEC-derived peaks (Peaks 1.I and 1.III) originating from AEC Peak 1 (Supplementary Figure S4A and S5A). The concentration of U at the maximum of Peak 1.III was approximately 1.3-fold higher than that measured at the edges of the same peak (Supplementary Figure S5). Accordingly, all spots with an intensity ratio of 1.3 between the maximum and each of the two flanking fractions were considered as candidate protein spots. For Peak 1.I, this ratio was 2.2 on the left and 1.13 on the right with respect to the top of the peak (Supplementary Figure S4). Each of these peaks was analyzed in two independent experiments. The candidate spots were selected according to two criteria. The first criterion, termed “standard”, was met when the spot intensity at the top of the U peak was greater than the intensities of the spots on either side, in both experiments. The second criterion, termed “strong”, was more stringent. It was met if the lowest intensity of the spot at the top of the U-peaks (between the two experiments) was higher than the higher intensity of the spots on either side of the U-peaks. Therefore, any spot meeting the strong criterion also satisfied the standard criterion, but the reverse was not necessarily true. Although this method may not reveal all UraBPs within a protein spot, it effectively identifies the most abundant candidates, providing a focused list for subsequent validation of U(VI)-binding capacity.

**Figure 2.**
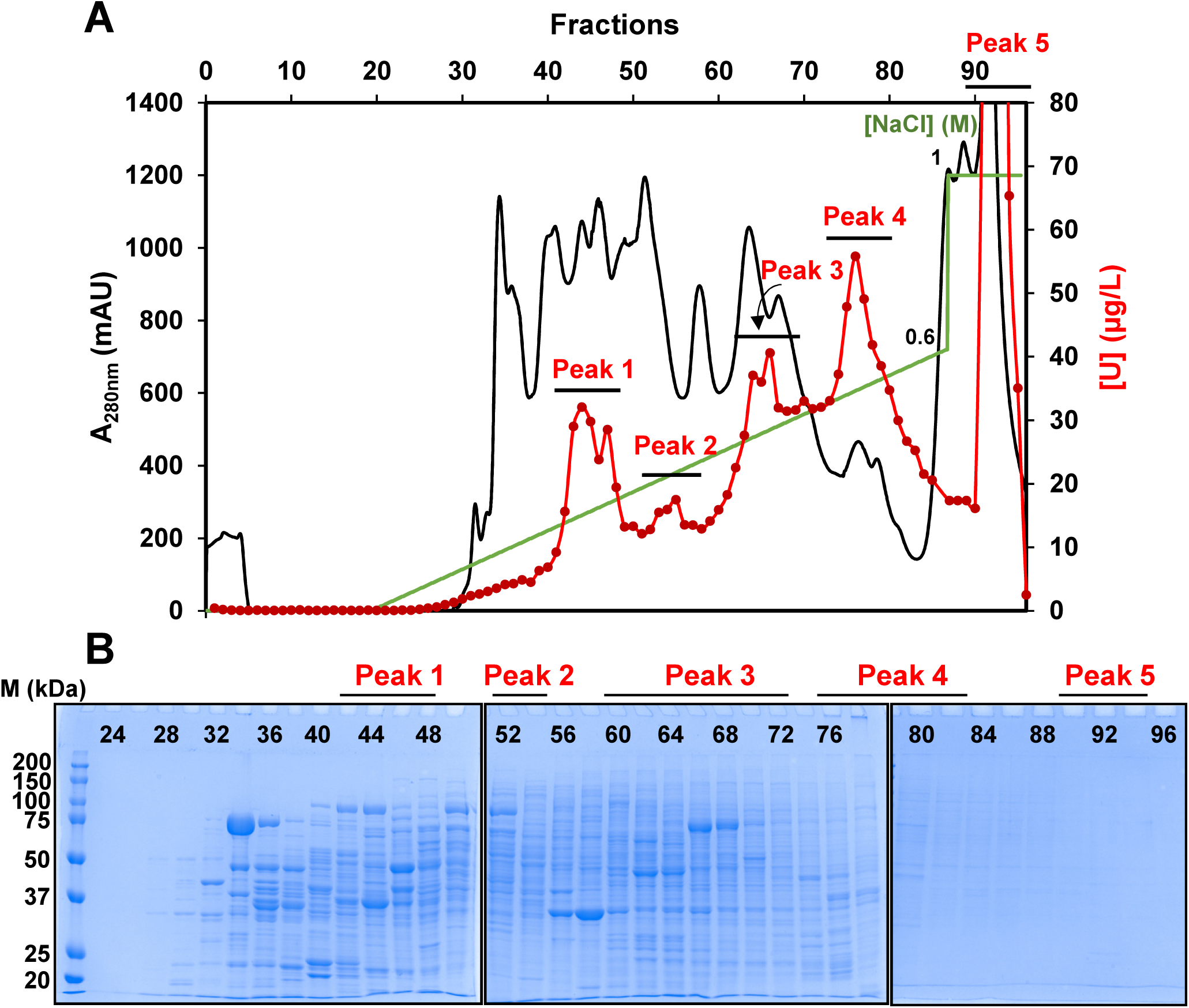
Fractionation by anion exchange chromatography of a soluble protein extract isolated from *A. thaliana* cells challenged with uranyl nitrate. This represents the first step of strategy 1 for identification of UraBPs. **A.** Three grams of soluble proteins from 200 g of *A. thaliana* cells, treated with 50 µM uranyl nitrate for 24 h, were separated on a Q-Sepharose HP column by aliquots of 300-400 mg of proteins (details are given in the Materials and Methods section). The protein profile is shown in black, U quantified in each eluted fraction by ICP-MS is shown in red, and the linear salt gradient from 0 to 0.6 M NaCl is in green. The elution revealed five distinct U peaks (1 to 5). This graph is representative of the same experiment performed 9 times independently. **B.** SDS-PAGE analyses of fractions eluted from the column, after Coomassie Blue staining. One fraction out of two, from 24 to 96, have been analyzed (10 µl/well).

In the case of Peak 1-I, fraction 35 corresponding to the top of the U peak and fractions 32 and 37 corresponding to the left and right sides of the peak, respectively (Supplementary Figure S4A) were separated on 2D SDS-PAGE (Supplementary Figure S4B). Sixteen candidate UraBP spots were selected, nine of which met the strong criterion and seven only the standard criterion (Supplementary Figure S4C). Proteins present in these spots were identified by nLC-MS/MS. These proteins and their functions are listed in Supplementary Table S2, along with some of their physicochemical properties, including the proportion of amino acids typically involved in U(VI) interactions (Glu, Asp, Tyr, His) and phosphorylated residues (pSer, pThr, pTyr), which enhance U(VI) binding to Asp and Glu [22, 23, 38, 39, 49–56]. The table also highlights any known relationship between these proteins and metals, including metal-binding ability and/or regulation by metals. For example, three of these proteins have known phosphorylated residues and ten are known to bind metals or are regulated by metal stress (Supplementary Table S2). One of them, the phosphoenolate carboxykinase (Uniprot id: Q9T074), was found in three protein spots, one of which has an enrichment rate greater than 3 (Spot 4) (Supplementary Figure S4C). This protein is multi phosphorylated (5 known sites) and is involved in the response to cadmium stress [57].

Similarly, fraction 55 (top of the peak) and fractions 53 and 58 (left and right sides of the U peak) were analyzed for Peak 1-III (Supplementary Figure S5A). The analysis of protein spot intensities highlighted 29 candidate spots (Supplementary Figure S5B), 22 of which met the strong criterion and 7 of which met only the standard criterion (Supplementary Figure S5C). A list of 23 candidate proteins was obtained (Supplementary Table S3). Six spots showed an average abundance change (“fold change”) with the top of the peak close to 2 (spots 6, 7, 8, 11, 12 and 13). Among them, spot 8 corresponds to the aminomethyltransferase T-protein of the glycine cleavage system (Uniprot ID: O65396), known to respond to cadmium stress. Another protein (spot 11) corresponds to an isocitrate dehydrogenase (Uniprot ID: Q8LG77), which requires Mg^2+^ or Mn^2+^ ions as cofactor for its activity. Another candidate protein, lactoylglutathione lyase (Uniprot ID: Q8H0V3), also called glyoxylase, stands out because it is a zinc-binding protein with a binding site containing two Glu and one His, amino acids with affinity for U(VI). It is also involved in the detoxification of methylglyoxal, a compound produced by lipid peroxidation induced by oxidative stress [58]. Of note, the probable 3-hydroxyisobutyrate dehydrogenase-like 1 protein (Uniprot ID: Q9SZE1) was identified as a UraBP candidate in both the 1-I and 1-III peaks.

### 3.3. UraBP identification by Strategy 2: Towards the identification of the most affine proteins for U(VI) in cellulo

In strategy 2, 250 mg of proteins extracted from 13 g of U(VI)-treated cells (containing approximately 72 ng U/mg protein) were subjected to chromatography on a Bio-Gel HTP hydroxyapatite column as the initial purification step (Figure 1). Although the hydroxyapatite resin poses a risk of disrupting U(VI)-protein interactions due to phosphate strong affinity for U(VI), proteins with sufficiently high binding affinity can retain their association with U(VI) and are protected from precipitation by phosphate from the medium [45, 59]. Protein and U profiles are shown in Supplementary Figure S6. A significant part of U initially present in the protein extract was detected in the column exclusion volume. This fraction contains all the proteins not retained by the column as well as U(VI) weakly or non-specifically bound to proteins and precipitated by the phosphate present in the column equilibration buffer. A second U peak was observed eluting at approximately 150 mM phosphate, representing around 5% of the total uranium and 24% of the proteins from the original extract. This fraction was likely to contain UraBPs in a stable complex with U. Fractions from this peak were pooled and subsequently separated using a Superdex 200 SEC column. The resulting U profile was composed of five distinct peaks (Supplementary Figure S6) containing 0.2; 3.5; 9.3; 18 and 0.3 mg of protein, respectively. Notably, the fifth peak, comprising low molecular weight proteins (< 20 kDa), emerged as a key area of interest due to its low protein complexity and potential enrichment in high-affinity UraBPs (Supplementary Figure S6, Figure 3). The proteins present in the five peaks were separated in a final chromatographic step on a high-resolution Q-Sepharose column. The most striking result was the fractionation of peak 5 from the Superdex 200 column. Most of the U from this peak eluted as a narrow peak in fraction 26 (F26) of the Q-Sepharose column (Supplementary Figure S6, Figure 3). This fraction contained only five detectable bands, as determined by SDS-PAGE and silver nitrate staining. Proteins present in this fraction as well as in peak 5 of the Superdex 200 column were identified by nLC-MS/MS. A list of 23 candidate proteins was retained after sorting the protein as function of their size (<20 kDa) and ranking according to their relative abundance (“Specific Spectral counts”) (Supplementary Table S4). Some of these proteins, such as the monothiol glutaredoxin-S12 (Uniprot ID: Q8LBS4), the probable calcium-binding protein CML13 (Uniprot ID: Q94AZ4), and the 16 kDa phloem protein 1 (Uniprot ID: Q9M2T2), are metalloproteins. Some proteins are multi-phosphorylated, such as the nucleoside diphosphate kinase 1 (Uniprot ID: P39207), or are enriched in Glu and Asp residues, like the probable calcium-binding protein CML13 (Uniprot ID: Q94AZ4) and 16 kDa phloem protein 1 (Uniprot ID: Q9M2T2). However, the glycine-rich RNA-binding proteins 7 and 8 (GRP7 and GRP8) were at the top of the list. In addition to being the most abundant proteins in the fraction, both paralogs are multi-phosphorylated (12 and 8 known sites, respectively). In addition, GRP7 was recently identified as a protein with *in vitro* affinity for U(VI), after affinity capture of Arabidopsis leaf and root protein extracts, on U(VI)-immobilized matrix [26]. GRP7 is a well-characterized small soluble RNA-binding protein containing a globular N-terminal RNA recognition motif (RRM) and a C-terminal glycine-rich intrinsically disordered region (IDR) [60]. It regulates the expression of numerous genes at the post-transcriptional level, including its own transcripts and the GRP8 transcripts [61], by controlling pre-mRNA splicing and/or translation [62]. Target genes are involved in various biotic [63] and abiotic stresses response [62, 64–67], including metal stress [68].

**Figure 3.**
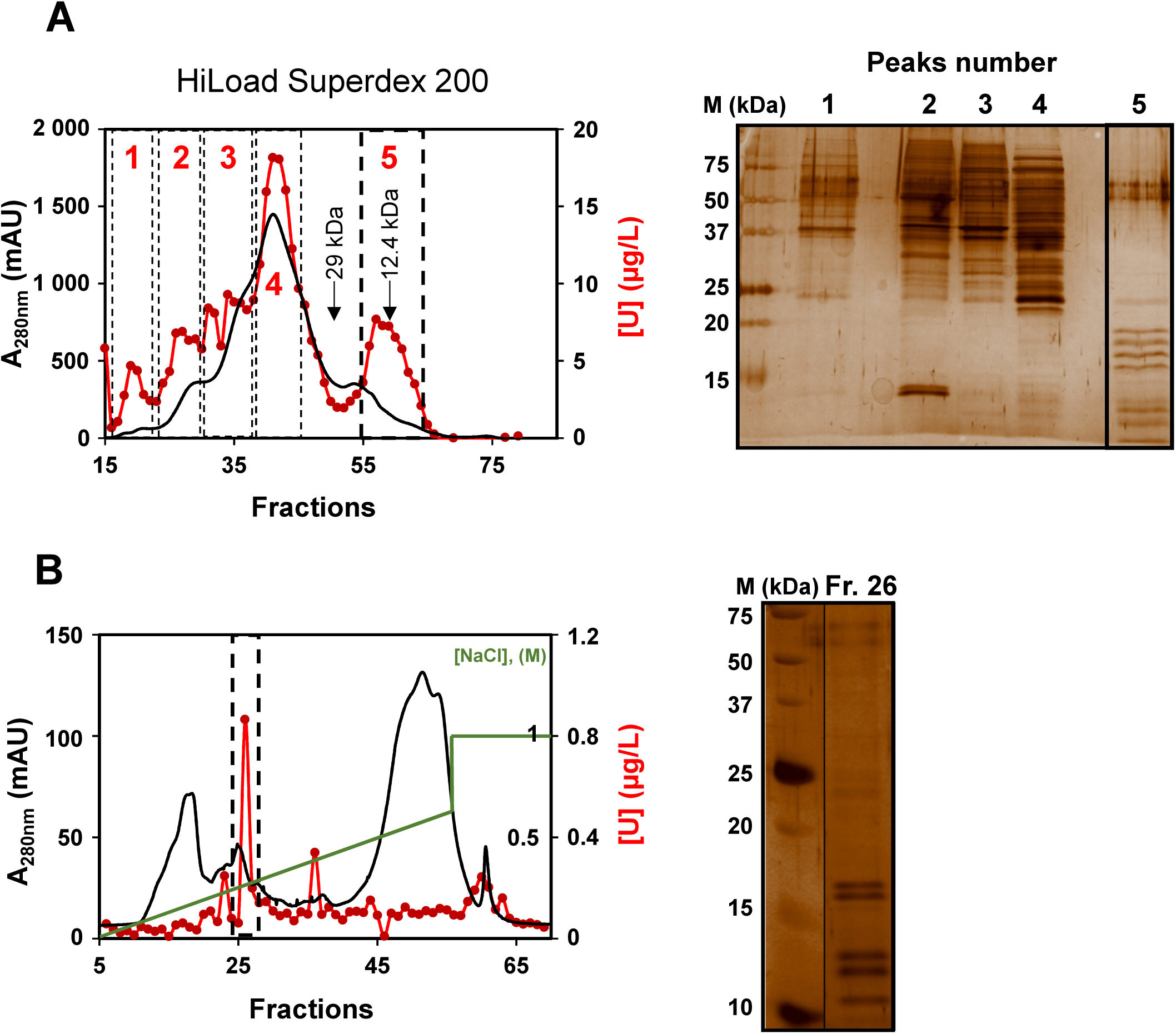
Strategy 2 for the identification of UraBPs from *A. thaliana* cells challenged with uranyl nitrate. **A.** The major U peak eluted from a Bio-gel HTP hydroxyapatite column (Supplementary Figure S6) was further chromatographed onto an HiLoad Superdex 200 column. The protein profile is shown in black and U, quantified in each eluted fraction by ICP-MS, is in red. Elution revealed five distinct U peaks (1 to 5). The elution of molecular weight markers (carbonic anhydrase, 29 kDa; cytochrome C, 12.4 kDa) in an independent chromatography is indicated by arrows. SDS-PAGE analysis of U peaks (0.2-1 µg protein from pooled peak fractions/ well), after silver staining is shown on the right. **B.** Uranium peak 5 from the SEC column was chromatographed on a Q-Sepharose HP column. The protein profile is in black, U profile in red, and the linear salt gradient from 0 to 0.6 M NaCl in green. The elution revealed one major U peak, eluted in fraction 26. SDS-PAGE analysis of U peak in fraction 26 (Fr. 26, 10µl), after silver staining is shown on the right.

In light of all these considerations, we focused on GRP7 to achieve a more detailed characterization.

### 3.4. Recombinant full-length GRP7 protein binds 2 U(VI) ions per monomer through intramolecular interactions

To confirm the ability of *A. thaliana* GRP7 to bind U(VI) and to further characterize its metal-binding properties, we cloned the cDNA encoding the full-length GRP7 protein by RT-PCR and produced the native (without any tag) recombinant protein in the *E. coli* Rosetta2 (DE3) strain. We used a three-step procedure including two-successive chromatographic steps to purify the recombinant GRP7 protein to near homogeneity (Supplementary Figure S7A). Because we observed that the GRP7 solutions became reversibly turbid at low temperatures (10°C or less) and low salt concentrations (<250 mM NaCl), 300 mM NaCl was included in the SEC media in the course of protein purification and in all subsequent buffers used in this study. Indeed, GRP7 is known to undergo phase separation under heat or cold stress *in vitro* and *in vivo*, and this phenomenon is dependent on protein and salt concentration [62]. Using 2 L of bacterial culture, we were able to purify up to 12 mg of recombinant GRP7. Finally, the recombinant GRP7 protein behaved as an apparent ≍16.5 kDa globular protein by size exclusion chromatography (SEC) on a Superdex 200 Increase 10/300 GL column, as calculated from a calibration curve obtained by measuring elution volumes of calibration proteins, indicating that the protein is monomeric in solution (Supplementary Figure S7B).

To analyze the U(VI)-binding capacity of recombinant GRP7 *in vitro* and to determine the stoichiometry of U(VI) complexation, we used the arsenazo III assay to determine the U(VI) content in U(VI)-GRP7 complexes formed after incubation of 15 μM GRP7-buffered samples with 200 μM uranyl nitrate (∼13 U(VI) equivalents). Separation of complexes from the unbound metal was achieved by centrifugation on Sephadex MicroSpin G-25 columns, as described (Vallet et al., 2023) (Supplementary Figure S8A). Under these conditions, the data showed a ligand binding of 2.14 ± 0.25 U(VI) ions per GRP7 monomer, consistent with the presence of 2 U(VI) binding sites on the recombinant protein (Supplementary Figure S8B). None of the other metal ions tested, including Ca(II), Fe(II) and Fe(III), Zn(II), Ni(II), Cd(II) and Pb(II), bound significantly to recombinant GRP7 (<0.2 metal-equivalent binding; not shown). Finally, U(VI) binding did not affect the protein elution profile on SEC, indicating that U(VI) binding was intramolecular and did not alter its oligomerization state (Supplementary Figure 7B).

### 3.5. Solution NMR investigation of GRP7

To gain insights into the *in vitro* molecular interaction of U(VI) with GRP7, we performed solution NMR spectroscopy on ^13^C/^15^N labeled protein samples. All NMR data sets were recorded, with purified GRP7 (typically 100 µM) in Tris-HCl 20 mM pH 7.5, NaCl 0.3 M, with magnetic field strengths of either 700 MHz or 850 MHz ^1^H frequency, at 27°C. A ^1^H-^15^N correlation spectrum of full length GRP7 is shown in Figure 4A. A first observation from this spectrum is that the number of cross peaks is much lower than the number of amide groups in the backbone of this 176-residue protein. Site-specific NMR assignments were obtained from a set of 3D HNC-type Best-TROSY experiments [41]. The assigned chemical shifts revealed that only the N-terminal part of GRP7 (residues 5-86) gives rise to observable NMR signals under our experimental conditions, while the glycine-rich C-terminal part is hardly observable. This finding is in-line with predictions of disorder/order scores (Figure 4B) that indicate a high disorder propensity in the C-terminal half of the protein. The absence of notable NMR intensity detected for this part of the protein indicates that the GRP7 C-terminal domain is not forming a highly flexible polypeptide chain, but rather a molten globular structure with interconversion of multiple conformations on the milliseconds time scale. NMR chemical shifts also provide information on the secondary structural elements present in the N-terminal part of GRP7 (Figure 4C; [69]). The identified β–α–β–β–α–β topology is typical of RNA recognition motifs (RRM), in agreement with another recent NMR study on GRP7 [60] and the AlphaFold structural model. We further confirmed that the RRM fold is independent of the glycine-rich C-terminal domain. For this aim, we prepared a truncated GRP7 construct, named GRP72, which only comprises residues 1-90. The ^1^H-^15^N spectrum of GRP72 overlaps almost perfectly with the spectrum of full-length GRP7, indicating no changes in domain structure and conformational dynamics upon removing of the C-terminal part (Figure 4A).

**Figure 4.**
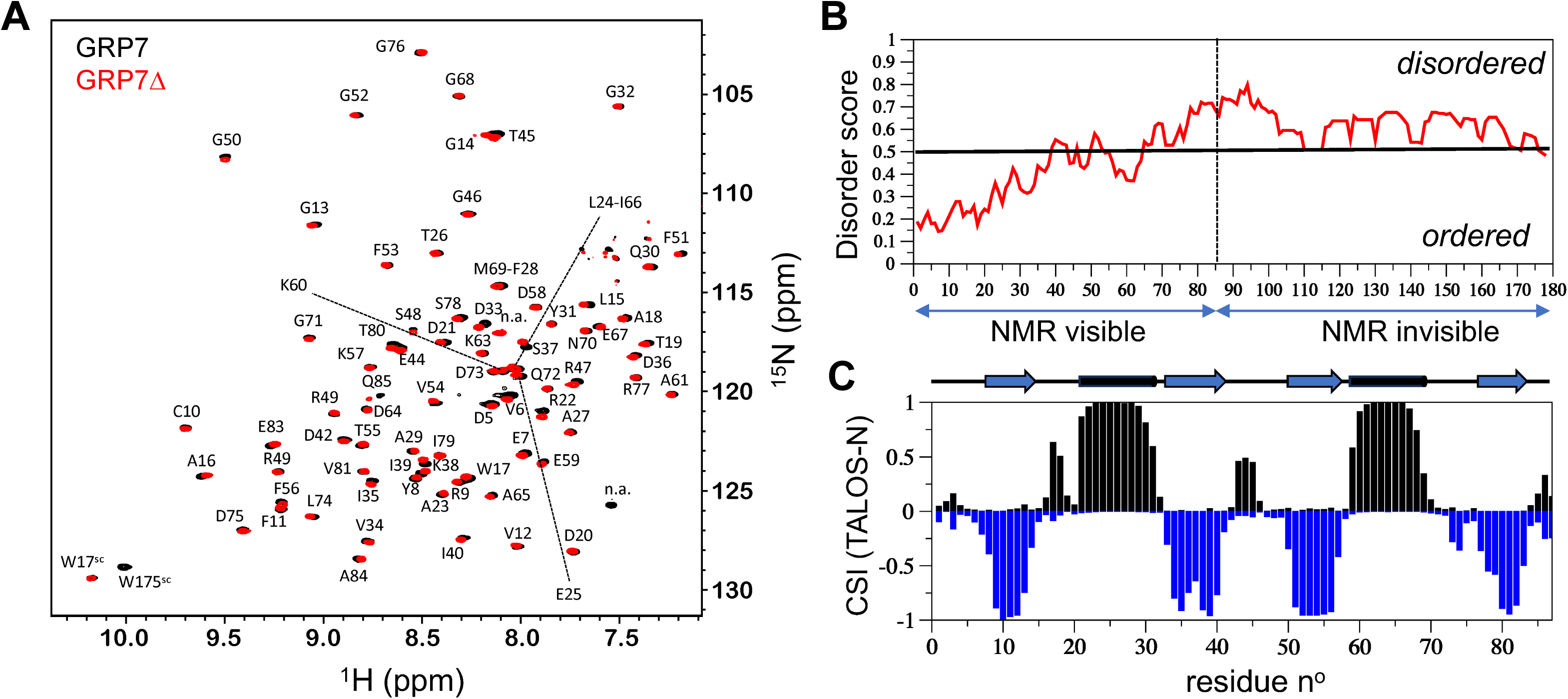
NMR investigation of the apo GRP7 recombinant proteins. **A.** Overlay of the ^1^H-^15^N correlation spectra of the full-length GRP7 (black contours) and GRP72 (red contours) recorded at 27°C and 700 MHz ^1^H frequency. Peaks are assigned by amino-acid type and residue number. Only the protein part from residues 5 to 86 give rise to NMR signals in these spectra. **B.** Conformational disorder score of the full-length GRP7 computed with the IUPred software and plotted as a function of the protein sequence. **C.** Secondary structural propensities of the N-terminal part of full-length GRP7 (5-86) computed a chemical shift index (CSI) from NMR chemical shifts using TALOS-N [69]. The helical score (positive) is plotted in black while the β-strand score (negative) is plotted in blue. Secondary elements as identified from this CSI are plotted on top.

We then investigated the interaction of full-length GRP7 with U(VI). Comparing ^1^H-^15^N correlation spectra recorded for the apo GRP7 and a 1:2 GRP7:U(VI) sample showed no significant changes in peak positions (Figure 5A). However, a quantitative analysis of spectra revealed a significant increase in peak intensity upon U(VI) binding for a few residues (amide groups Asp5, Trp17, Ser48, Gln85, and Ser86) (Figures 5A and 5B). In addition, small peak shifts observed in aliphatic side chain ^1^H-^13^C correlation spectra allowed to identify one more residue (Ala2) affected by the interaction with U(VI). Interestingly, all NMR-identified nuclear sites that pick up the presence of U(VI) are located either at the N- and C-terminal ends of the RRM domain, or in loop regions connecting secondary structural elements. The position of these residues on the AlphaFold2 structural model highlights two spatially distinct regions as potential U(VI) binding sites, named U1 and U2 (Figure 5C). Taking into account the binding stoichiometry determined by our biochemical assay, we hypothesized that each of these binding sites is able to coordinate a single U(VI) cation. Our NMR data did not show any indication that the glycine-rich C-terminal part is implicated in the interaction with U(VI). This was further confirmed by NMR data recorded on the truncated GRP72 showing similar spectral changes as observed for full-length GRP7 (Supplementary Figure S9), and by biochemical data showing almost identical U(VI) binding stoichiometry to GRP72 compared to full-length GRP7 (Supplementary Figure S8B). Finally, we addressed whether U(VI) coordination involves only a monomer of GRP7 or several GRP7 molecules forming oligomeric complexes. For this, we performed NMR-based translational diffusion experiments (Figure 5D). No change in the apparent diffusion constant was observed upon the addition of 2 molar equivalents of U(VI), demonstrating that GRP7 remains monomeric upon U(VI) binding, consistent with SEC analysis (Supplementary Figure 7B).

**Figure 5.**
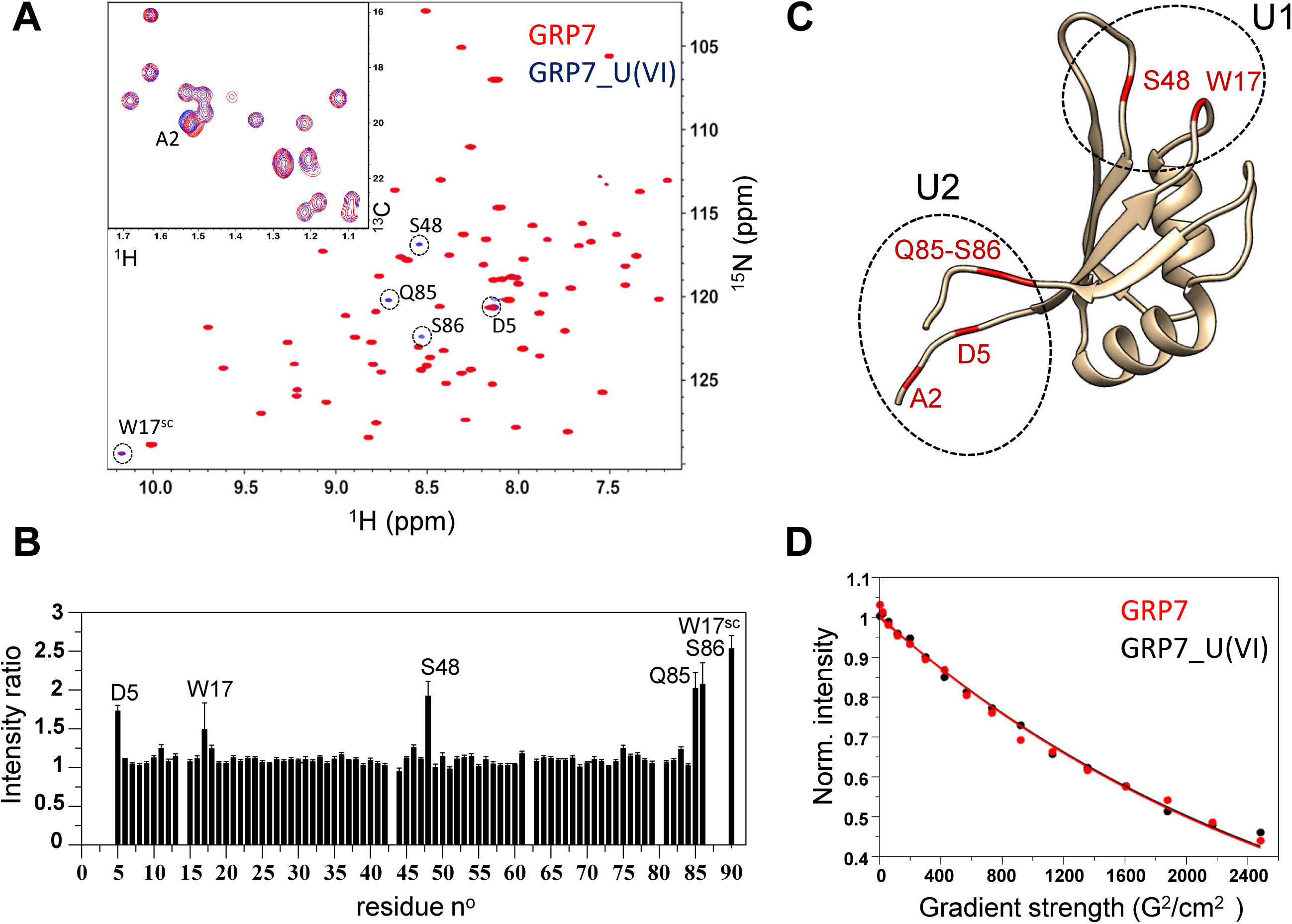
NMR characterization of the interaction between the full-length GRP7 and uranyl. **A.** Superposition of the amide ^1^H-^15^N spectra recorded for the apo GRP7 (red) and a 1:2 GRP7:U mixture (black). GRP7 residues showing significant changes in peak intensity upon uranyl addition are highlighted by dashed circles and annotated. A small region of the methyl ^1^H-^13^C spectra, recorded on the same samples is shown as an insert. The NMR peak of residue A2 shows a small frequency shift between the two samples. **B.** NMR peak intensity ratios (GRP7-U complex/apo GRP7) computed for individual amide sites and plotted as a function of protein sequence. While for most residues this ration is close to 1 (no change in conformation and dynamics), some residues (annotated) experience an increase in peak intensity upon complex formation. **C.** Structural model of GRP7. The model was obtained with AlphaFold v2.0. All residues showing increased amide signal intensities or peak shifts of side chain resonances are located at the N- and C-terminal end of the structured domain, as well as in flexible loop regions. **D.** Translational diffusion NMR measurements (DOSY) show no difference in apparent particle size between the apo GRP7 (red) and the GRP7-U complex (black).

### 3.6. Characterization of U(VI)-binding residues of GRP7

To confirm and refine the putative U(VI)-binding sites on GRP7 and to identify the residues that coordinate the U(VI) ions, we first aligned the GRP7 and GRP8 RRM domain sequences with those of orthologous human RNA-binding proteins, previously found to bind U(VI) *in cellulo* [48]. In particular, we focused on identifying conserved Glu and Asp residues located on the vicinity of U(VI)-sensitive residues observed in NMR studies (Figure 6A). Indeed, oxygen from the carboxyl side chain of these amino acids is a hard Lewis base known to be one of the main functional groups for high affinity U(VI) binding to proteins, through equatorial coordination of up to six amino acid ligands, perpendicular to the U-O-U axis [22, 56]. This analysis identified Asp42 and Glu44, on one hand, and Asp5 and Glu7, on the other hand, as putative U(VI)-binding residues within or in the vicinity of U1 and U2 putative binding sites, respectively (Figure 6B). In order to confirm that these residues are indeed involved in U(VI) coordination, we have produced a set of mutants of GRP72 modified on both U1 (Asp42Ala/Glu44Ala) and U2 (Asp5Ala/Glu7Ala) sites and checked their ability to bind U(VI) (Figure 7). Our results show that mutations of each individual site significantly reduced U(VI) binding, while mutations of both sites almost completely abolished U(VI) binding. Mutation of the Asp42 and Glu44 residues to Ala reduced the U(VI) binding stoichiometry to GRP72 by 2-fold, suggesting that these mutations disrupt U1 binding site. Similarly, mutating the Asp5 and Glu7 residues to Ala also significantly reduced U(VI) binding, although to a lower extent, indicating that additional residues contribute to U(VI) coordination at the U2 site. However, all our attempts to produce recombinant GRP72 with triple mutations targeting either the U1 (Asp42Ala/Glu44Ala/Ser48Ala) or the U2 (Asp5Ala/Glu7Ala/Ser86Ala) sites were unsuccessful.

**Figure 6.**
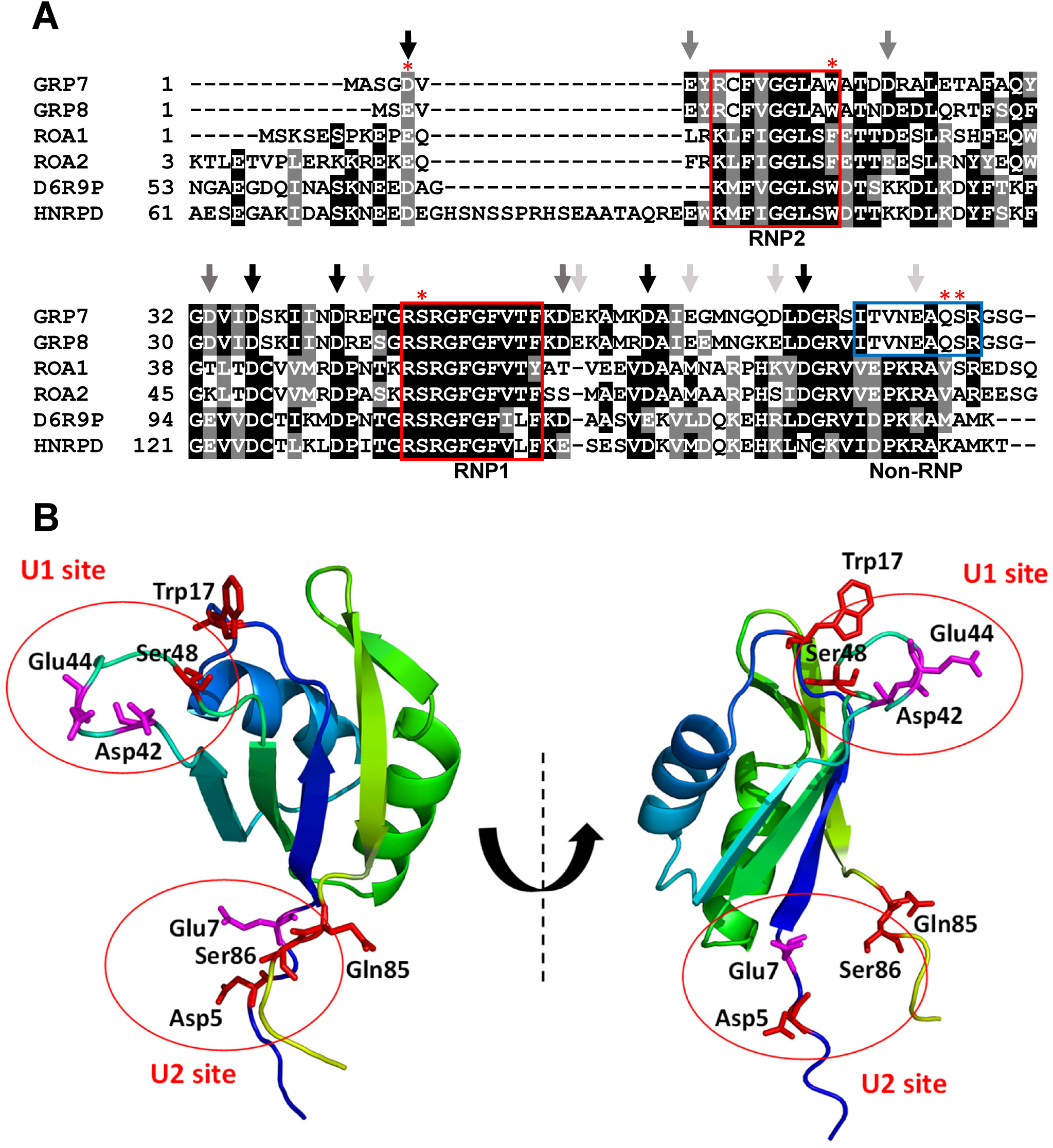
Identification of putative uranyl-binding residues on recombinant *A. thaliana* GRP7. **A.** Alignment of the *A. thaliana* GRP7 and GRP8 RNA-recognition motif (RRM) domains with the uranyl-affine human orthologous RNA-binding proteins, ROA1, ROA2, D6R9P and HNRPD. Multiple sequence alignment was performed using ClustalW. Conserved Glu and Asp residues are indicated by black (strong consensus), grey (medium consensus) and light grey (Arabidopsis proteins only) arrows. Ribonucleoprotein (RNP) motifs within the RNA recognition motif domain are highlighted with red boxes. The non-canonical RNA interaction domain extension (Non-RNP) for Arabidopsis proteins is in the blue box. GRP7 residues sensitive to U(VI) in NMR are indicated by red stars. **B.** Structural model of GRP72. The model was obtained using Alphafold v2.0 and the image was produced using PyMOL (DeLano Scientific, SanCarlos, CA, USA) as a ribbon model colored in the “chainbows” mode. Position of NMR sensitive residues to U(VI) and neighboring Glu and Asp residues are displayed as red and purple sticks, respectively.

**Figure 7.**
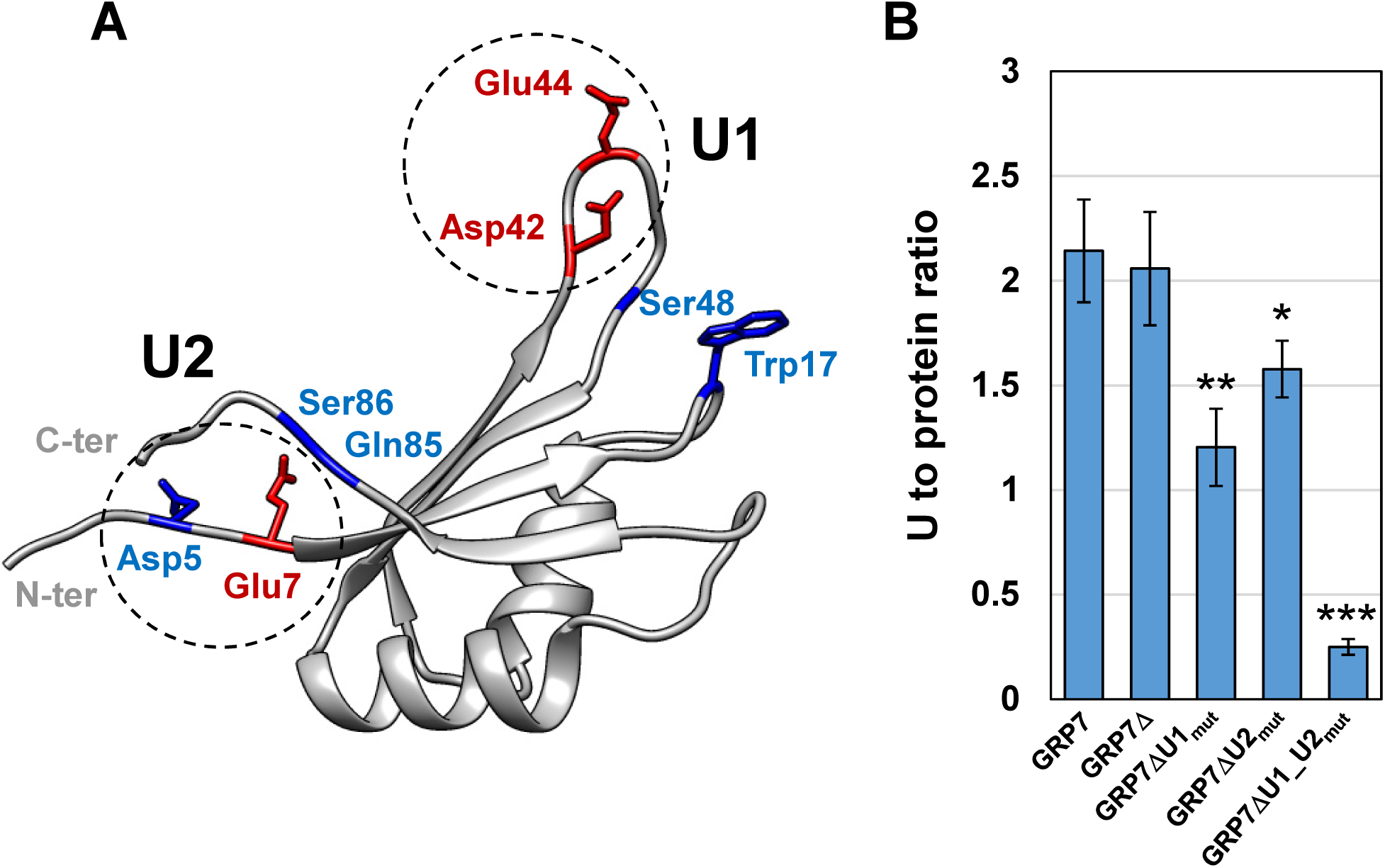
Effect of mutations in the GRP7Δ uranyl-binding sites U1 and U2 on U(VI) binding stoichiometry. **A.** Position of NMR sensitive residues to U(VI) (represented as blue sticks) and neighboring Glu and Asp residues (red sticks) within the putative U1 and U2 GRP72 uranyl-binding sites. Residues Asp42 and Glu44 for site U1, and Asp5 and Glu7 for site U2 have been mutated to Ala in GRP72 mutants. **B.** Determination of U(VI) binding stoichiometry in GRP72 mutants. Uranium was determined using the arsenazo III assay in filtrates (containing protein-U(VI) complexes) obtained by centrifugal size exclusion chromatography of GRP7, GRP72 and GRP72 mutants solutions (15 µM), incubated with 200 µM U(VI), in the presence of 150 µM iminodiacetic acid. See Material and Methods for experimental details. Values are mean ± SD of 4 independent measurements (Supplementary Figure S8); * p-value ≤ 0.05; ** p-value ≤ 0.01; *** p-value ≤ 0.001; Dunnett test.

### 3.7. Nucleic acid binding to GRP7 interferes with U(VI) binding

GRP7 has been shown to play pivotal roles in the post-transcriptional regulation of gene expression. By binding to specific RNA sequences, GRP7 can modulate RNA splicing, stability, and translation, thereby fine-tuning the expression of stress-responsive genes [70]. This regulation is particularly important under stress conditions, where precise control of gene expression can determine the extent of plant adaptation and survival. In a recent work, Lewinski et al. [60] identified GRP7 RNA targets from individual-nucleotide resolution UV cross-linking and immunoprecipitation (iCLIP) data [71] and determined a conserved RNA motif enriched in uridine residues at the GRP7 RRM binding sites. Using NMR titrations, they optimized the 7-mer 5’ AGUUUCA RNA ligand comprising the GRP7 consensus binding motif determined from iCLIP binding sites. This oligonucleotide binds specifically and with high affinity to the conserved ribonucleoprotein (RNP) motifs RNP1 and RNP2 and to a newly identified non-consensus motif (Non-RNP) from the RRM domain [60]. The corresponding single-stranded DNA oligonucleotide 5’ AGTTTCA 3’ binds to the GRP7 RRM domain with similar affinity and specificity. Interestingly, some of the GRP7 residues involved in sensing the binding of U(VI), as identified by NMR or by site-directed mutagenesis, such as Trp17, Asp42, Ser48, Gln85 and Ser86, are either identical or located close to those that interact with the consensus RNA or DNA oligonucleotide within the RRM domain (Figure 6A; [60]). This suggests a possible interference between oligonucleotides and U(VI) binding to the GRP7 RRM core domain. To test this hypothesis, we pre-incubated GRP72 with increasing concentration of the single-stranded DNA oligonucleotide 5’ AGTTTCA 3’ before the addition of 60 µM U(VI). U(VI) binding was then assessed by SEC of protein complexes and Arsenazo III assay. The results presented in Figure 8 show that binding of the consensus oligonucleotide to GRP72 prevented U(VI) binding in a dose-dependent manner. In marked contrast, incubation of the protein with the single-stranded DNA oligonucleotide 5’ AAAAAAA 3’, which does not interact with GRP7 [60], had no or very little effect on U(VI) binding.

**Figure 8.**
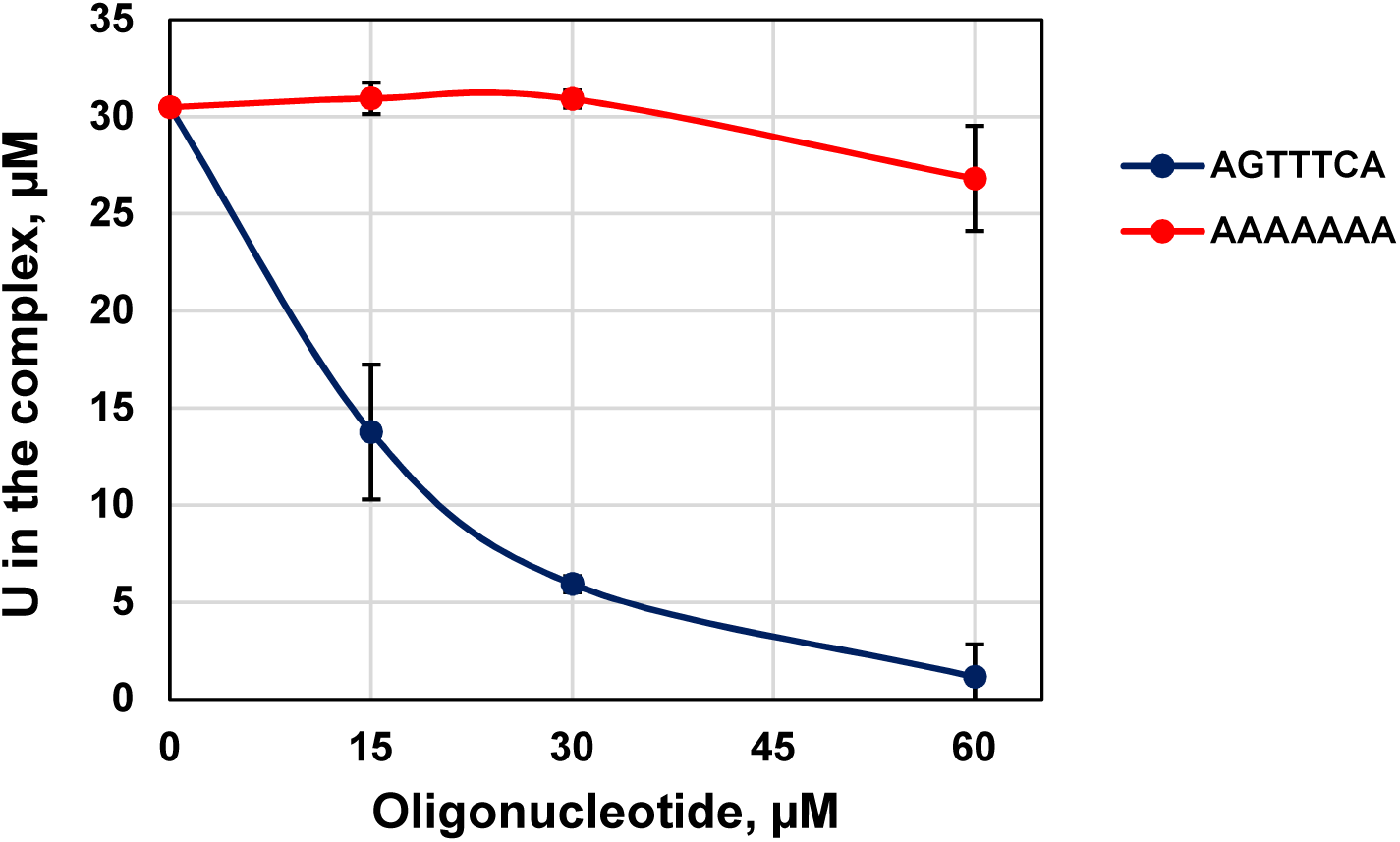
Effect of oligonucleotide binding to the GRP7 RRM domain on protein interaction with U(VI). Uranium was determined using the arsenazo III assay in filtrates (containing protein-U(VI) complexes) obtained by centrifugal size exclusion chromatography of GRP72 solutions (15 µM) preincubated for 10 min with increasing concentrations of the single-stranded DNA oligonucleotide 5’ AGTTTCA 3’ or 5’ AAAAAAA 3’, and subsequently incubated for 30 min with 60 µM U(VI) and 150 µM iminodiacetic acid. See Material and Methods for experimental details. Values are mean ± SD of 3 independent measurements.

## 4- Discussion

Uranium, particularly in its hexavalent form U(VI), has been shown to induce various toxic effects across all living species. Once internalized by cells, U(VI) interacts predominantly with cellular biomolecules, especially proteins. Previous studies have elucidated the mechanisms by which U(VI) penetrates living cells, notably highlighting the involvement of iron(III) and calcium(II) uptake pathways in facilitating U entry into eukaryotic organisms [17, 18]. Building on this knowledge, the present study aimed to identify soluble protein targets of U(VI) in plant cells. Proteins, as key biomolecules orchestrating numerous cellular processes and metabolic pathways, are central to the mechanisms of toxicity induced by non-essential metals, including U. Numerous proteins have been reported to interact with U(VI), most of which in human systems. These interactions can provoke structural alterations in proteins, as exemplified by modifications observed in osteopontin [47, 72] and fetuin A [50, 72]. Such interactions are sometimes helpful to elucidate the mechanisms of U(VI) toxicity. For instance, U(VI) disrupts the cytb5-cytc complex, triggering apoptosis [73, 74]. Moreover, U(VI) can substitute essential biological metals such as iron and calcium in metalloproteins, disrupting the functions of proteins like transferrin, the primary iron transporter in blood [75], and calmodulin [38, 76]. Uranyl also interferes with DNA-binding transcription factors by targeting zinc finger motifs [77] and exhibits photocatalytic properties leading to protein and DNA cleavage [78, 79]. Investigations into U(VI)-binding proteins *in vivo* have been conducted in various organisms, including bacteria [45], rats [80], crayfish [81], and humans [48, 56, 82–84]. However, despite extensive studies on the toxic effects of U(VI) in plants, no *in vivo* U(VI) protein targets have been identified to date. To bridge this gap, we employed a metalloproteomics approach combining advanced protein separation techniques, designed to optimally preserve U(VI)-protein interactions, with robust protein and metal identification methods (nLC-MS/MS, coupled or not with two-dimensional electrophoresis and ICP-MS). These methodologies were applied to protein extracts from *A. thaliana* cell suspension cultures exposed to uranyl nitrate.

The use of *A. thaliana* cell suspension cultures allowed for the collection of substantial protein quantities while bypassing the histological complexity inherent to higher plants. Additionally, these poorly differentiated cells exhibit relatively uniform protein expression across the *A. thaliana* proteome [27], reducing the likelihood that highly abundant proteins with low U(VI) affinity would obscure the detection of UraBPs with high affinity but low expression levels. For example, albumin was long considered the principal U(VI) target in blood serum [85]. Subsequent findings revealed that albumin binds only 7% of serum U(VI), whereas fetuin A, despite its lower abundance, binds over 80% due to its significantly higher affinity (apparent Kd of 30 nM compared to 17 μM for albumin) [50]. Thus, the utilization of plant cell cultures grown under heterotrophic conditions effectively mitigates this masking effect, enhancing the identification of specific high-affinity U(VI)-binding proteins.

In our pioneering experiments, we exposed cells to a low-phosphate medium [27], as phosphate is known to significantly limit U(VI) uptake [5, 86, 87]. The complexation of U(VI) with phosphate results in the formation of highly insoluble compounds [88], leading to U(VI) precipitation in the medium or its accumulation on plant cell walls. Consequently, low amounts of soluble U(VI) were detected within the cells under these conditions. To enhance U(VI) uptake, we reduced cell density to increase the U(VI)/biomass ratio and used a phosphate-free medium during exposure to U(VI) nitrate. These adjustments did not compromise cell viability [3] or growth over the 24-hour exposure period (Supplementary Figure S1). Under these optimized conditions, the proportion of soluble intracellular U increased from less than 2% (in low-phosphate conditions) to nearly 20% of total U, with approximately half of this soluble fraction bound to proteins (Supplementary Figure S2). These conditions proved suitable for isolating and identifying UraBPs.

A straightforward and reliable method for identifying UraBPs involves a chromatographic approach utilizing modified IMAC. This method exploits the cation exchange properties of aminophosphonate groups to selectively bind U(VI) [24]. This strategy has previously enabled the identification of numerous U(VI)-binding proteins in human kidney and neuronal cells *in vitro* [25, 48]. Applying this technique, we recently identified 38 U(VI)-affine proteins in protein extracts from *A. thaliana* roots and leaves. Among these, a protein designated UraBP25 has been further characterized [26]. UraBP25 is notably rich in glutamate residues, which are recognized as preferential binding sites for U(VI), and it exhibits nanomolar-range affinity for U(VI). However, while this method effectively identifies proteins with *in vitro* affinity for U(VI), it does not confirm their actual interaction with U(VI) *in planta* or in cultured cells exposed to U(VI).

The identification of intracellular protein targets of U(VI) is the central objective of this study, providing essential insights into the cellular mechanisms underlying U(VI) toxicity and tolerance. Two principal methodologies are commonly employed for this purpose: chromatographic and electrophoretic techniques, often used in combination to maximize efficiency. Chromatographic separation facilitates protein fractionation and enrichment of UraBPs. Complementarily, two-dimensional gel electrophoresis under native conditions, followed by laser ablation coupled with ICP-MS, enables the precise localization of U and subsequent identification of UraBPs through mass spectrometry (nLC-MS/MS) [80, 81, 89–91]. In their exploration of bacterial metalloproteomes, Cvetkovic et al. [45] demonstrated the efficacy of column chromatography for enriching proteins bound to essential metals such as iron, zinc, and calcium. However, no similar enrichment was observed for toxic metals like U. This discrepancy could be attributed to the nonspecific binding of U(VI) to numerous proteins and the progressive dissociation of U(VI) from proteins with genuine affinity during separation steps. Notably, Cvetkovic et al. [45] utilized a hydroxyapatite column composed of a calcium phosphate matrix and phosphate buffer for elution. Given phosphate’s high affinity for U(VI), this setup likely promoted the precipitation of nonspecifically or weakly protein-bound U(VI) [59]. Considering the strengths and limitations of prior methodologies, we implemented two complementary chromatographic strategies. Strategy 1 focuses on preserving U(VI)-protein interactions to enable the comprehensive identification of UraBPs, irrespective of their binding affinity. In contrast, Strategy 2, which begins with an hydroxyapatite chromatography, selectively isolates proteins with the highest affinity for U(VI) (Figure 1). This dual approach balances exhaustive detection with targeted identification, enhancing our understanding of U(VI)-protein interactions.

The two UraBP identification strategies developed in this study led to the discovery of 57 potential protein candidates, 36 identified through Strategy 1 and 23 through Strategy 2. Notably, the putative 3-hydroxyisobutyrate dehydrogenase-like 1 (Uniprot ID: Q9SZ21) was detected in both the peak 1.I and 1.III fractions using Strategy 1, while the desiccation-related protein At2g46140 (Uniprot ID: O82355) was identified by both strategies. Candidate proteins were classified based on several criteria: the presence of at least two confirmed phosphorylation sites (multi-phosphorylation), metal-binding capability or involvement in metal stress responses, and an overrepresentation of amino acid residues with potential U(VI) affinity (Supplementary Tables S2, S3, and S4). Among proteins fulfilling multiple criteria, the iron-storage proteins ferritin-1 (Uniprot ID: Q39101) and ferritin-3 (Uniprot ID: Q9LYN2) were identified through Strategy 1 (Supplementary Table S2). Ferritins are universal intracellular proteins that store iron in its oxidized form and regulate iron homeostasis by controlling iron release [92]. Interestingly, metalloproteomic analyses in the hyperthermophilic archaeon *Pyrococcus furiosus* exposed to U(VI) identified ferritin as a major in vivo U(VI) target [45]. Similarly, ferritin was shown to bind U in the U-bioaccumulating crayfish *Procambarus clarkii* exposed to U(VI) contamination [91]. These findings suggest that U(VI), like other actinides, mimics iron chemistry [93] and can displace iron *in vivo*. This implicates ferritin as both a potential target of U(VI) toxicity and a player in U detoxification across diverse species. Moreover, since *A. thaliana* ferritin-1 and ferritin-3 are localized in chloroplasts, their interaction with U(VI) would suggest that U can penetrate chloroplasts. Another compelling UraBP candidate identified through Strategy 1 is a lactoylglutathione lyase (Uniprot ID: Q8H0V3), which naturally binds zinc via a coordination site formed by two Glu, one His, and one Gln residue [94]. This binding site could plausibly accommodate U(VI), forming a pentagonal bipyramidal coordination complex with U axial positions occupied by oxygen atoms [22, 82]. Among the 57 UraBP candidates identified in this study, particular attention was given to glycine-rich RNA-binding protein 7 (GRP7; Uniprot ID: Q03250), which emerged as the most enriched UraBP candidate, alongside its paralog GRP8 (Uniprot ID: Q03251), through metalloproteomics Strategy 2 (Supplementary Table S4). GRP7 had previously been reported to exhibit high-affinity binding to U(VI) *in vitro*, as demonstrated by a metalloproteomics study using modified IMAC affinity chromatography [26]. Additionally, a recent investigation into the urano-proteome of neuronal cells identified six heterogeneous nuclear ribonucleoproteins (hnRNPs) as potential UraBPs [48]. Given that these hnRNPs share homology with plant GRP7 and GRP8, this further reinforced the decision to prioritize their characterization. GRP7 and GRP8 belong to class IV of plant glycine-rich proteins (GRPs), also referred to as RNA-binding GRPs [70]. This subclass is involved in plant responses to biotic and abiotic stress, primarily through post-transcriptional gene regulation [95]. Structurally, these proteins feature a highly conserved N-terminal RRM of 80–90 amino acids, adjacent to a C-terminal glycine-rich intrinsically disordered region (IDR). This IDR is believed to mediate interactions with various components involved in RNA processing and protein transport between the nucleus and the cytosol [96]. The RRM domain is also critical for RNA processing by GRPs, enabling phase separation of proteins in complex with RNA and promoting the assembly of stress granules under stress conditions [62, 97].

Here, using a combination of biochemical and structural analyses, we demonstrated that purified recombinant GRP7 binds U(VI) *in vitro* at two distinct intramolecular sites within its RRM domain (Supplementary Figure S8; Figure 5). This mode of interaction contrasts with U(VI) binding to PCaP1 from A. thaliana observed in a previous work, where U(VI) induced oligomerization by bridging monomer interfaces at distinct sites in both the structured N-terminal domain and the Glu-rich flexible C-terminal region [26]. High-resolution NMR spectroscopy of GRP7 revealed a significant increase in peak intensities in ^1^H-^15^N correlation spectra for specific residues at opposite edges of the RRM domain upon addition of a twofold excess of U(VI) to the apo-protein (Figure 5). This suggests direct or proximal interaction with U(VI). A similar approach identified Glu34 and Asp38 as U(VI)-binding residues in the cyanobacterial protein SmtA [49], though in that case, interactions were detected through chemical shift changes, a common but not exclusive signature of protein-ligand binding [98]. Site-directed mutagenesis confirmed the involvement of four key residues in GRP7 RRM domain U(VI) binding: Asp42 and Glu44 within the U1 site, and Asp5 and Glu7 within the U2 site (Figure 7). Mutations in the U1 site completely abolished U(VI) binding, while those in the U2 site only partially reduced binding, implying that additional residues may contribute to U(VI) coordination. The two oxo groups of the U(VI) cation facilitate the coordination of up to six ligands in its equatorial plane [22, 56], suggesting a complex binding network. Interestingly, Ser85, sensitive to U(VI) exposure in recombinant GRP7 in NMR studies, has been reported to be phosphorylated *in vivo* (Supplementary Table S4). Given the well-established role of phosphorylation in enhancing U(VI) binding affinity [38, 53, 99, 100], Ser85 may participate in U(VI) interaction within the U2 site, further strengthening binding affinity *in planta*.

Our data indicate that U(VI) binding to GRP7 is exclusive to the RRM domain, with no involvement of the glycine-rich domain (Supplementary Figures S8 and S9; Figure 7). However, the glycine-rich domain undergoes extensive phosphorylation (up to 12 sites in GRP7 and 8 in GRP8) (Supplementary Table S4; [62, 101]) suggesting that in native conditions, this domain may also participate in U(VI) binding. This aligns with the efficient purification of native GRP7 under stringent conditions [26] and the isolation of GRP7:U(VI) and GRP8:U(VI) complexes *via in cellulo* metalloproteomics Strategy 2 in this study.

Beyond its well-established roles in circadian clock regulation [102–104], stress tolerance to cold, drought, salinity and temperature fluctuations [62, 65, 105], pathogen immunity [63, 106], and floral transition [107], all intrinsically linked to its RNA-binding capacity [101, 107–109], GRP7 has also emerged as a key regulator in heavy metal stress responses in *A. thaliana*. Notably, GRP7 expression is modulated under Cd [57], Zn [110], Pb, Ni [68], and U [15] exposure. However, unlike U, these metals do not appear to bind GRP7, at least *in vitro*. Recent findings by Kim et al. [68] unveiled GRP7 novel roles in mediating *A. thaliana* tolerance to Ni and Pb. GRP7 transcript levels increased in response to Ni but decreased under Pb exposure. Moreover, GRP7 was shown to regulate the mRNA stability of target genes involved in heavy metal chelation and antioxidant defense, influencing the plant capacity to accumulate and tolerate Ni and Pb. Our study further demonstrates that U(VI) binding occurs specifically within the RRM domain of GRP7, targeting regions overlapping or adjacent to the RNP1, RNP2, and non-RNP motifs (Figure 6). These binding events interfere with the interaction of the optimized GRP7 target oligonucleotide. (Figure 8), suggesting a compelling hypothesis: U(VI) interaction with GRP7, and by extension GRP8, *in planta* could impair the RNA-binding and processing functions of these multifaceted proteins. Such interference may significantly affect plant responses to U-induced stress. Exploring this mechanism could uncover new strategies for enhancing plant resilience to U exposure.

## 5- Conclusions

The identification and characterization of UraBPs in plants is a pivotal step toward unraveling the molecular mechanisms underlying plant responses and adaptations to U(VI) stress. These proteins could act as direct targets of U toxicity or play critical roles in metal detoxification and tolerance. In this study, we developed and optimized robust metalloproteomics strategies to isolate and identify UraBP candidates from *A. thaliana* cultured cells. Our objective was to pinpoint proteins that interact with U(VI) in plant cells. These approaches led to the identification of 57 candidate UraBPs. Among these, GRP7 emerged as a particularly compelling candidate, previously identified through an independent metalloproteomics study targeting *in vitro* U(VI)-affine proteins [26]. Further characterization of GRP7, using integrated biochemical and structural analyses, confirmed its ability to bind U(VI) at two distinct intramolecular sites within its RRM domain. This binding was challenged by nucleotide target binding, suggesting that GRP7 could be a direct molecular target of U toxicity in plants. These findings not only validate our methodological approach but also provide critical insight into how U(VI) may interfere with essential RNA-binding proteins, potentially impairing key post-transcriptional regulatory processes. This knowledge opens new avenues for exploring plant defense mechanisms against U stress and developing crops with enhanced metal tolerance, contributing to sustainable agriculture and phytoremediation strategies.

### Environmental implications

The characterization of uranyl-binding proteins (UraBPs) in plants, identified through *in vitro* and *in vivo* metalloproteomics, provides valuable insights into the mechanisms of uranium (U) chemical toxicity and detoxification. This fundamental knowledge is a prerequisite for the sustainable management of U contamination in polluted soils and throughout the food chain. Moreover, this understanding could drive the development of innovative biotechnological solutions for the remediation of contaminated water sources and the creation of biosensors. Such advancements may be achieved by optimizing U(VI)-binding sites on proteins or engineering derived mimetic peptides [22, 39, 55, 111].

### Data Availability

All data supporting the findings of this study are included in this published article (and its Supplementary Information files), or are available from the corresponding authors upon reasonable request. NMR chemical shifts of GRP7 have been deposited with the BMRB (http://www.bmrb.wisc.edu/) under accession number 52967. Proteomics data have been deposited to the ProteomeXchange Consortium via the PRIDE partner repository [112] with the dataset identifier PXD056391 and Project DOI: 10.6019/PXD056391 (Strategy 1); and dataset identifier PXD058046 and Project DOI: 10.6019/PXD058046 (Strategy 2).

### Author contributions: CRediT

**Benoit H. Revel:** Conceptualization, Data curation, Formal analysis, Investigation, Methodology. **Adrien Favier:** Conceptualization, Data curation, Formal analysis, Investigation, Methodology, Validation. **Jacqueline Martin-Laffon:** Formal analysis, Investigation. **Alicia Vallet:** Conceptualization, Data curation, Formal analysis, Investigation, Methodology, Validation. **Jonathan Przybyla-Toscano:** Data curation, Writing – review and editing. **Sabine Brugière:** Investigation. **Yohann Couté:** Conceptualization, Data curation, Formal analysis, Investigation, Methodology, Writing – review and editing. **Hélène Diemer:** Investigation. **Sarah Cianférani:** Conceptualization, Data curation, Formal analysis, Investigation, Methodology. **Thierry Rabilloud:** Conceptualization, Data curation, Formal analysis, Investigation, Methodology, Writing – review and editing. **Jacques Bourguignon:** Conceptualization, Data curation, Funding acquisition, Methodology, Project administration, Supervision, Validation, Writing – review and editing. **Bernhard Brutscher:** Conceptualization, Data curation, Formal analysis, Methodology, Validation, Writing – review and editing. **Stéphane Ravanel:** Conceptualization, Data curation, Funding acquisition, Methodology, Supervision, Validation, Writing – review and editing. **Claude Alban:** Conceptualization, Data curation, Formal analysis, Funding acquisition, Investigation, Methodology, Project administration, Supervision, Validation, Writing – original draft.

## Acknowledgments

Anne-Marie Boisson is kindly acknowledged for her technical assistance in *Arabidopsis thaliana* cell cultures.

This work was funded by grants from the Agence Nationale de la Recherche (ANR-17-CE34-0007, GreenU project; ANR-17-EURE-0003, CBH-EUR-GS; ANR-10-INBS-08-3, the French Proteomics Infrastructure, (ProFI)). The PhD fellowship to B.R. was funded by the CEA (CFR Grant).

This work used the high-field NMR platforms of the Grenoble Instruct-ERIC center (ISBG; UAR 3518 CNRS-CEA-UGA-EMBL) within the Grenoble Partnership for Structural Biology (PSB), supported by FRISBI (ANR-10-INBS-0005-02) and CBH-EUR-GS (ANR-17-EURE-0003).

Financial support from IR INFRANALYTICS FR2054 for conducting the research is also gratefully acknowledged.

## Appendix A. Supporting information

Supplementary data associated with this article can be found in the online version.

## Notes

### Competing Interest Statement

The authors have declared no competing interest.

