## Supplementary data for "Identification of uranyl-binding proteins in *Arabidopsis thaliana* cells exposed to uranium: Insights from a metalloproteomic analysis and characterization of Glycine-Rich RNA-binding Protein 7 (GRP7)"

### Supporting information

**Supplementary Table S1.** Synthetic oligonucleotides used in this study.
**Supplementary Table S2.** Identification of UraBP candidates - Strategy 1 (peak 1.I). *Available in a separate Excel file*
**Supplementary Table S3.** Identification of UraBP candidates - Strategy 1 (peak 1.III). *Available in a separate Excel file*
**Supplementary Table S4.** Identification of UraBP candidates - Strategy 2 (peak 5). *Available in a separate Excel file*
**Supplementary Figure S1.** Impact of uranium exposure on *A. thaliana* cultured cells growth.
**Supplementary Figure S2.** Uranium distribution in *A. thaliana* cells exposed to 50 µM uranyl nitrate in medium depleted with phosphate (NP medium).
**Supplementary Figure S3.** Second fractionation step of strategy 1 for identification of UraBPs.
**Supplementary Figure S4.** Differential analysis of putative UraBPs present in Peak 1.I by 2D-PAGE.
**Supplementary Figure S5.** Differential analysis of putative UraBPs present in Peak 1.III by 2D-PAGE.
**Supplementary Figure S6.** Fractionation of UraBPs according to strategy 2.
**Supplementary Figure S7.** Purification of recombinant GRP7 and determination of its oligomerization state.
**Supplementary Figure S8.** Determination of U(VI) binding stoichiometry to recombinant GRP7 protein variants.
**Supplementary Figure S9.** NMR characterization of the interaction between GRP7Δ and U(VI).

**Supplementary Table S1. Synthetic oligonucleotides used in this study.**

| Oligonucleotide name | Sequence (5’ → 3’) |
|---|---|
| **GRP7 cloning and expression** | |
| • NCoI-GRP7_5’ | GAGAGACCATGGCGTCCGGTGATGTTGAG |
| • SalI-GRP7_3’ | GAGAGTCGACTTACCATCCTCCACCACCACC |
| • SalI-GRP7Δ_3’ | GAGAGTCGACTTAACCGCTTCCTCGTGACTGAGC |
| **Mutagenesis** | |
| • GRP7_Site U1_For (D42A/E44A) | CAAGATCATTAACGCTCGTGCGACTGGAAGATCAA GGGGATTCGG |
| • GRP7_Site U1_Rev (D42A/E44A) | CCGAATCCCCTTGATCTTCCAGTCGCACGAGCGTT AATGATCTTG |
| • GRP7_Site U1’_For (D42A/E44A/S48A) | CAAGATCATTAACGCTCGTGCGACTGGAAGAGCAA GGGGATTCGG |
| • GRP7_Site U1’_Rev (D42A/E44A/S48A) | CCGAATCCCCTTGCTCTTCCAGTCGCACGAGCGTT AATGATCTTG |
| • GRP7_Site U2_For (D5A/E7A) | GGCGTCCGGTGCTGTTGCGTATCGGTGCTTC |
| • GRP7_Site U2_Rev (D5A/E7A) | GAAGCACCGATACGCAACAGCACCGGACGCC |
| • GRP7_Site U2’_For (S86A) | GTTAACGAGGCTCAGGCACGAGGAAGCGG |
| • GRP7_Site U2’_Rev (S86A) | CCGCTTCCTCGTGCCTGAGCCTCGTTAAC |
| **Competition oligo ssDNA *vs* U(VI)** | |
| • 7mer ssDNA | AGTTTCA |
| • 7mer polyA | AAAAAAA |

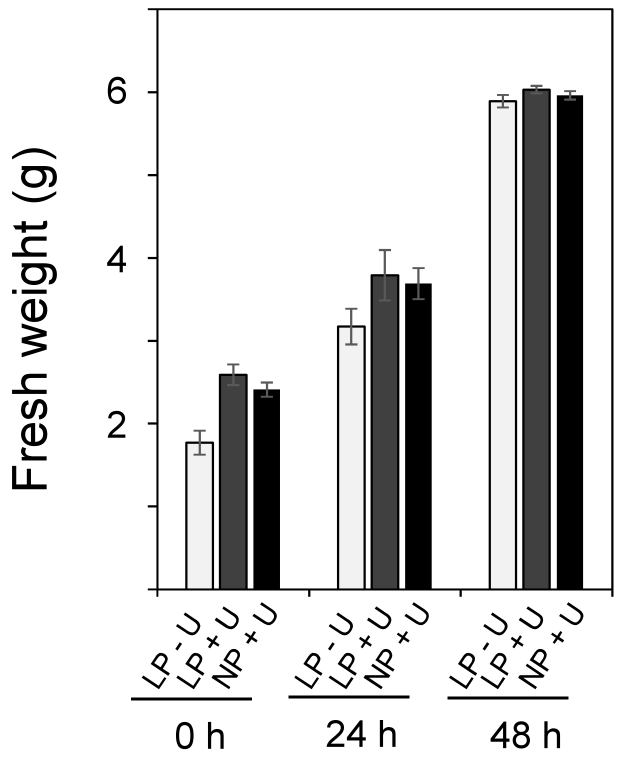


**Supplementary Figure S1. Impact of uranium on the growth of *A. thaliana* cultured cells.** Cells were grown in Murashige and Skoog medium to exponential phase (4 days after subculture) and then exposed to 50 µM uranyl nitrate in 100 ml medium with low phosphate (30 µM instead of 1.5 mM in regular medium) concentration (LP) or no phosphate (NP), for 0, 24h and 48h. After harvesting, the cells were washed with 10 mM sodium carbonate followed by water before fresh weight measurement. Data are mean ± SD of 3 independent measurements.

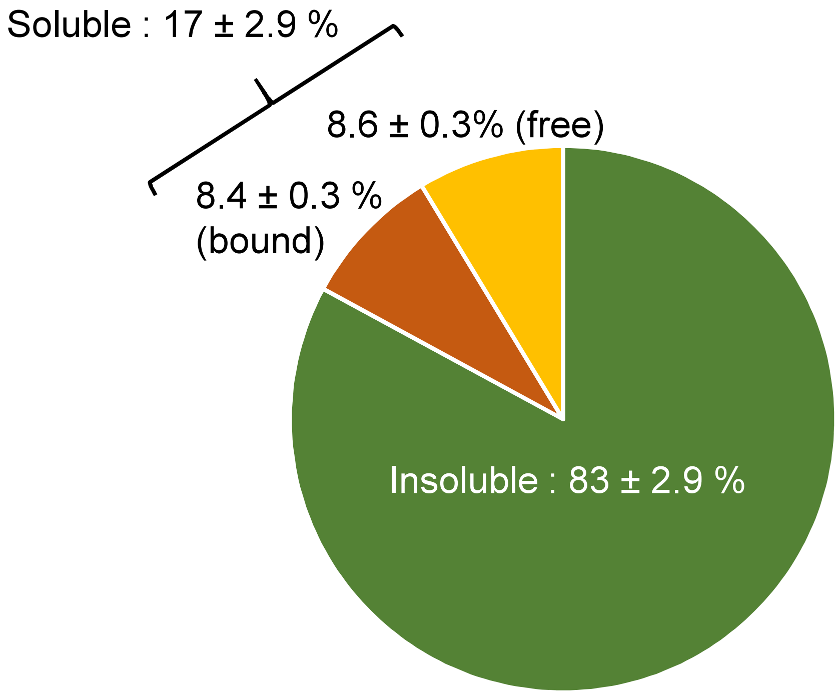


**Supplementary Figure S2. Uranium distribution in *A. thaliana* cells exposed to 50 µM uranyle nitrate in medium depleted with phosphate.** *A. thaliana* cell cultures concentrated to 2.5 g of fresh material per 100 ml of phosphate-depleted medium were exposed to 50 µM uranyl nitrate for 24 hours. After harvesting, the cells were washed with 10 mM sodium carbonate followed by water before cell disruption. The proportion of soluble and insoluble U was determined by ICP-MS by measuring U in the supernatant after cell lysis and centrifugation (see Materials and Methods). The soluble fraction consists of protein-bound U (orange) and "free" U (yellow). The latter two components are determined by ultrafiltration of the supernatant on a 3K filter. Uranium retained is bound to proteins larger than 3 kDa, whereas U found in the filtrate is considered to be 'free' or bound to small molecules (metabolites or peptides). The proportion of U in the insoluble form is shown in green and is determined by the difference between total U and that found in the soluble form. Data are mean ± SD of 5 independent measurements.

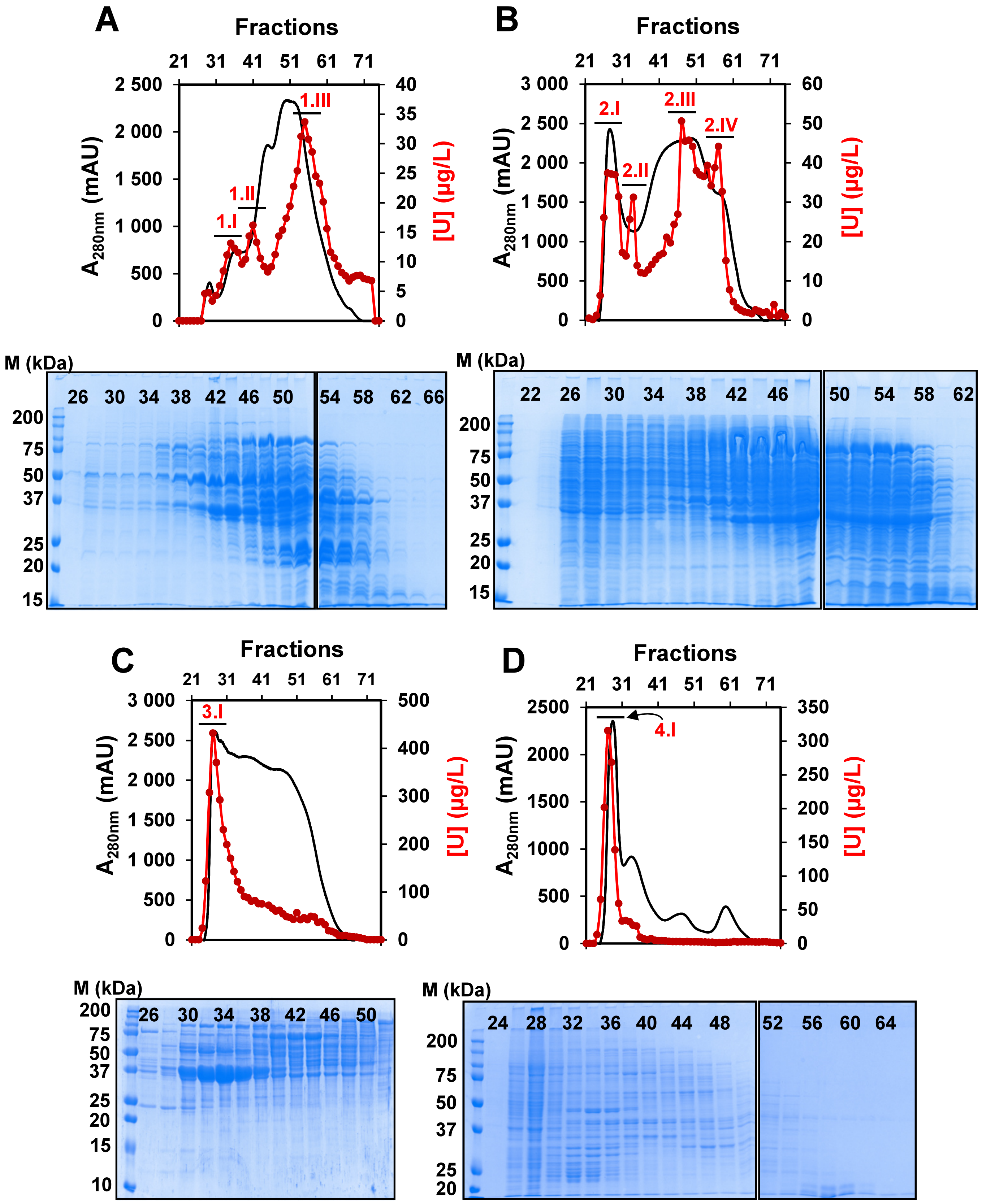


**Supplementary Figure S3. Second fractionation step of strategy 1 for the identification of UraBPs.** Proteins present in the U peaks 1 to 4 obtained after the first Q-Sepharose step (Figure 2) were separated by size exclusion chromatography on a Hiload Superdex 200 column. Peak 1 (**A**) gives rise to three U peaks (1.I to 1.III). Peak 2 (**B**) to four U peaks (2.I to 2.IV) and peak 3 (**C**) and 4 (**D**) to only one U peak (3.I and 4.I, respectively). The protein profiles are shown in black and U profiles in red. SDS-PAGE gels are shown under the corresponding chromatographic profiles. Aliquots (10 µl) of one eluted fraction out of two were analyzed. Fraction numbers are given.

**A**

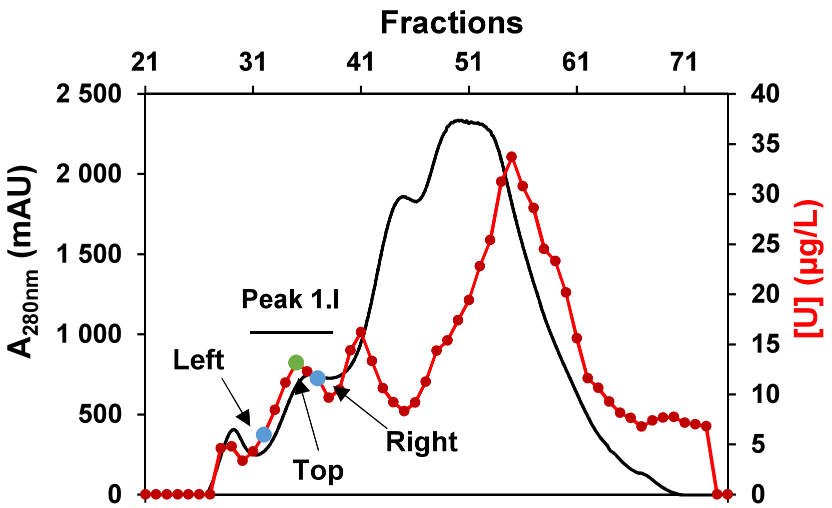


**B**

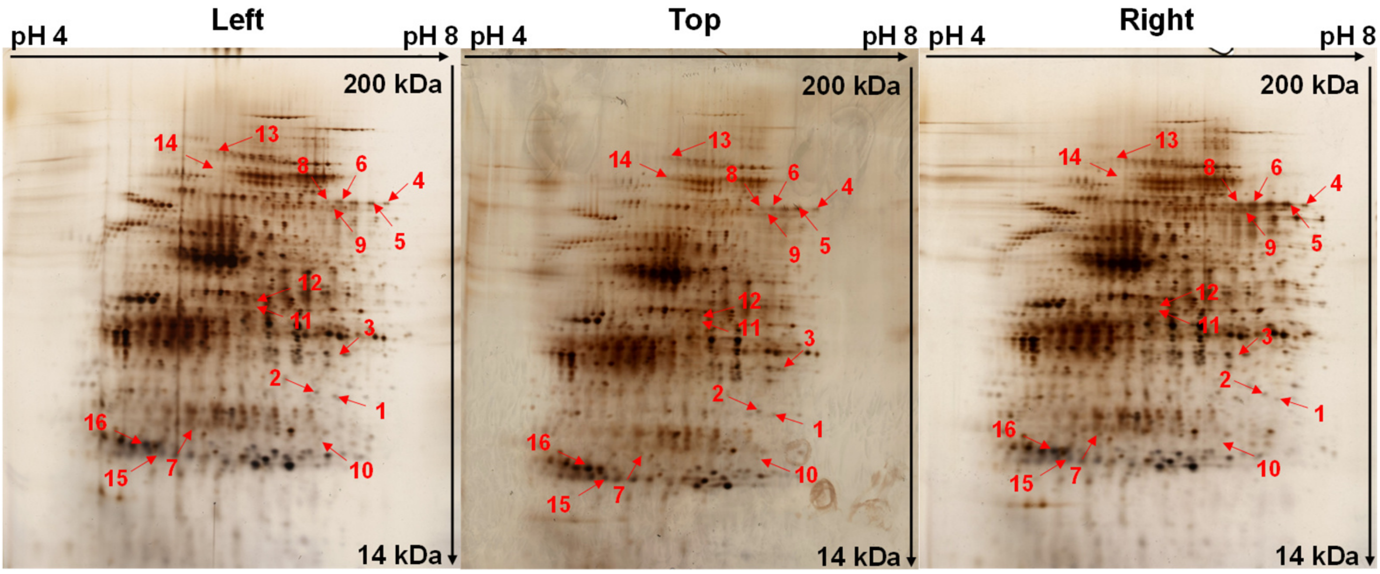


**C**

| Spot number | Intensities | | | Fold change | |
|---|---|---|---|---|---|
| | Left | Top | Right | Left | Right |
| **1** | **0.055** | **0.123** | **0.055** | **2.23** | **2.24** |
| **2** | **0.062** | **0.105** | **0.058** | **1.70** | **1.81** |
| **3** | **0.055** | **0.109** | **0.041** | **1.97** | **2.65** |
| **4** | **0.020** | **0.095** | **0.030** | **4.70** | **3.16** |
| **5** | **0.127** | **0.230** | **0.230** | **1.82** | **1.00** |
| **6** | **0.084** | **0.076** | **0.033** | **0.90** | **2.32** |
| **7** | **0.050** | **0.081** | **0.027** | **1.63** | **2.96** |
| **8** | **0.060** | **0.101** | **0.030** | **1.68** | **3.34** |
| **9** | **0.031** | **0.054** | **0.021** | **1.76** | **2.59** |
| 10 | 0.027 | 0.039 | 0.016 | 1.45 | 2.49 |
| 11 | 0.029 | 0.035 | 0.029 | 1.23 | 1.24 |
| 12 | 0.073 | 0.077 | 0.054 | 1.05 | 1.42 |
| 13 | 0.029 | 0.069 | 0.027 | 2.37 | 2.54 |
| 14 | 0.024 | 0.050 | 0.019 | 2.12 | 2.62 |
| 15 | 0.568 | 0.574 | 0.507 | 1.01 | 1.13 |
| 16 | 0.945 | 0.956 | 0.776 | 1.01 | 1.23 |

**Supp. Figure S4 (see next page for legend)**

**Supplementary Figure S4. Identification of putative UraBPs present in Peak 1.I by quantitative analysis of 2D-PAGE. A.** Superdex 200 column chromatography of peak 1 proteins from the Q-Sepharose column chromatography (Figure 2). **B.** 2D gel separation of fractions. **C.** Normalized relative intensity of the identified spot candidates. Three fractions of the peak 1.I (panel **A**) were analyzed on 2D gels stained with silver nitrate: the fraction at the top of the U peak and fractions of the left and right sides of the peak. Arrows show candidate protein spots whose relative intensity correlates with the U peak and which were analyzed and identified by mass spectrometry. The relative intensities of the protein spots were analyzed using the delta2D software and are shown in panel **C**, with the intensities of each candidate spot, selected according to either the strong criterion (bold) or the standard criterion (normal) (see the definition of these criteria in the main text), in the left, top and right fractions of the peak. Variations of spot intensities between the top of the peak and the left side (Fold change - left) and between the top of the peak and the right side (Fold change - right) are shown. The results presented are representative of two independent experiments.

**A**

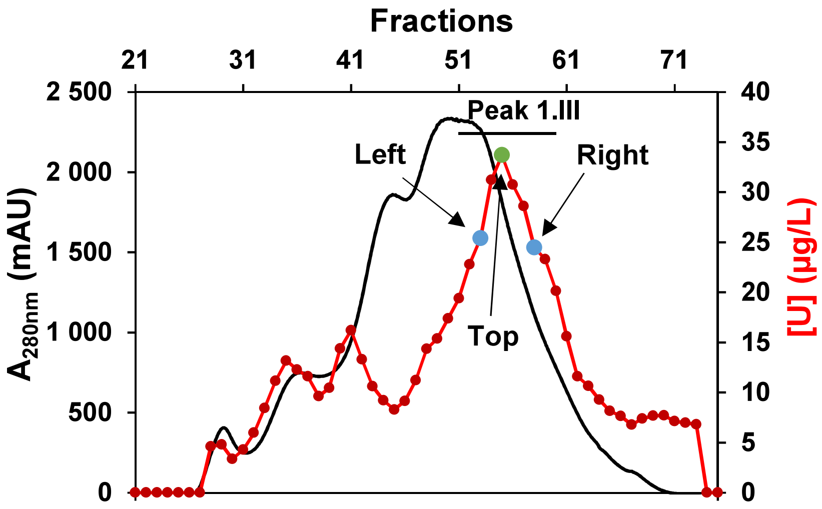


**B**

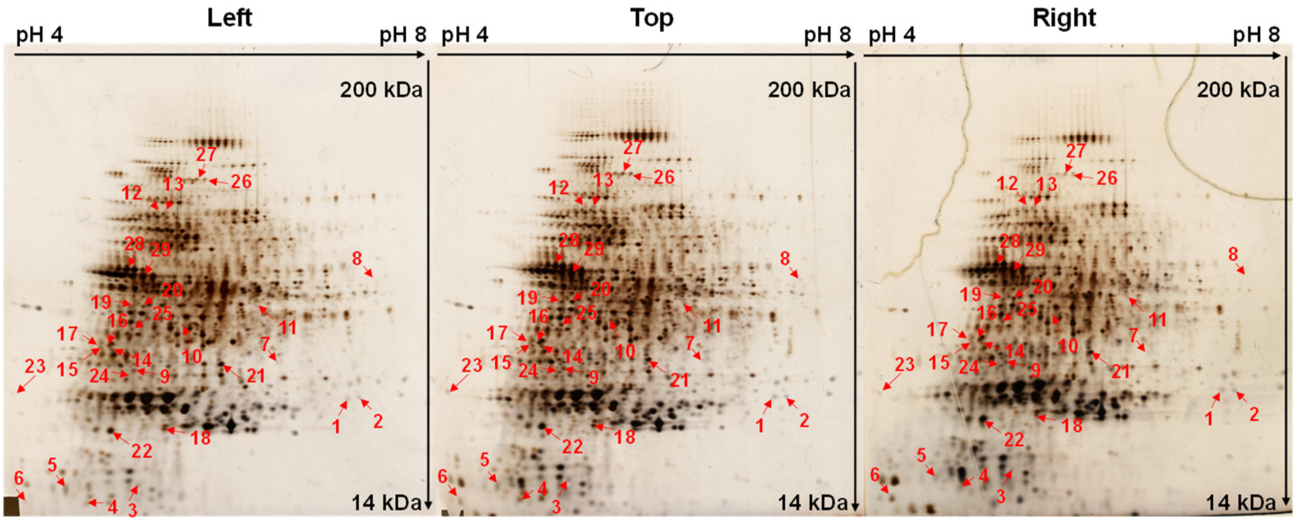


**C**

| Spot number | Intensities | | | Fold change | |
|---|---|---|---|---|---|
| | Left | Top | Right | Left | Right |
| **1** | **0.088** | **0.106** | **0.088** | **1.20** | **1.20** |
| **2** | **0.050** | **0.061** | **0.057** | **1.23** | **1.07** |
| **3** | **0.009** | **0.004** | **0.004** | **0.47** | **1.09** |
| **4** | **0.074** | **0.124** | **0.099** | **1.67** | **1.25** |
| **5** | **0.158** | **0.160** | **0.153** | **1.01** | **1.05** |
| **6** | **0.029** | **0.075** | **0.039** | **2.59** | **1.94** |
| **7** | **0.012** | **0.026** | **0.016** | **2.05** | **1.61** |
| **8** | **0.005** | **0.010** | **0.008** | **1.93** | **1.34** |
| **9** | **0.085** | **0.121** | **0.109** | **1.42** | **1.12** |
| **10** | **0.040** | **0.048** | **0.033** | **1.19** | **1.46** |
| **11** | **0.036** | **0.054** | **0.025** | **1.51** | **2.16** |
| **12** | **0.003** | **0.005** | **0.002** | **1.45** | **2.31** |
| **13** | **0.016** | **0.022** | **0.009** | **1.42** | **2.49** |
| **14** | **0.055** | **0.070** | **0.061** | **1.27** | **1.16** |
| **15** | **0.044** | **0.064** | **0.050** | **1.46** | **1.28** |
| **16** | **0.081** | **0.123** | **0.103** | **1.51** | **1.20** |
| **17** | **0.055** | **0.097** | **0.078** | **1.75** | **1.25** |
| **18** | **0.133** | **0.182** | **0.131** | **1.37** | **1.39** |
| **19** | **0.078** | **0.093** | **0.073** | **1.18** | **1.26** |
| **20** | **0.061** | **0.116** | **0.084** | **1.89** | **1.38** |
| **21** | **0.294** | **0.304** | **0.299** | **1.04** | **1.02** |
| **22** | **0.343** | **0.409** | **0.389** | **1.19** | **1.05** |
| 23 | 0.008 | 0.022 | 0.016 | 2.85 | 1.41 |
| 24 | 0.085 | 0.121 | 0.109 | 1.42 | 1.11 |
| 25 | 0.032 | 0.069 | 0.061 | 2.15 | 1.12 |
| 26 | 0.076 | 0.114 | 0.096 | 1.50 | 1.19 |
| 27 | 0.042 | 0.050 | 0.031 | 1.20 | 1.62 |
| 28 | 0.050 | 0.065 | 0.038 | 1.30 | 1.71 |
| 29 | 0.238 | 0.267 | 0.263 | 1.12 | 1.01 |

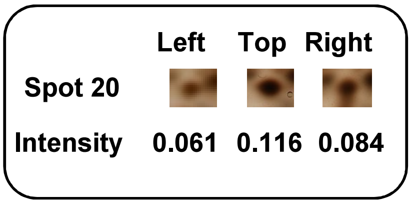


**Supp. Figure S5 (see next page for legend)**

**Supplementary Figure S5. Identification of putative UBP present in Peak 1.III by quantitative analysis by 2D-PAGE. A.** Superdex 200 column chromatography of peak 1 proteins from Q-Sepharose column chromatography (Figure 2) **B.** 2D gel separation of fractions. **C.** Normalized relative intensity of identified candidate spots. Three fractions of the peak 1.III (panel **A**) were analyzed on 2D gels stained with silver nitrate: the fraction at the top of the U peak and fractions of the left and right sides of the peak. Arrows show candidate protein spots whose relative intensity correlates with the U peak and which were analyzed and identified by mass spectrometry. The relative intensities of the protein spots were analyzed using the delta2D software and are shown in panel **C**, with the intensities of each candidate spot, selected according to either the strong criterion (bold) or the standard criterion (normal) (see the definition of these criteria in the main text), in the left, top and right fractions of the peak. Variations of spot intensities between the top of the peak and the left side (Fold change - left) and between the top of the peak and the right side (Fold change - right) are shown. The results presented are representative of two independent experiments. An example of spot analysis (spot 20) is given on the right.

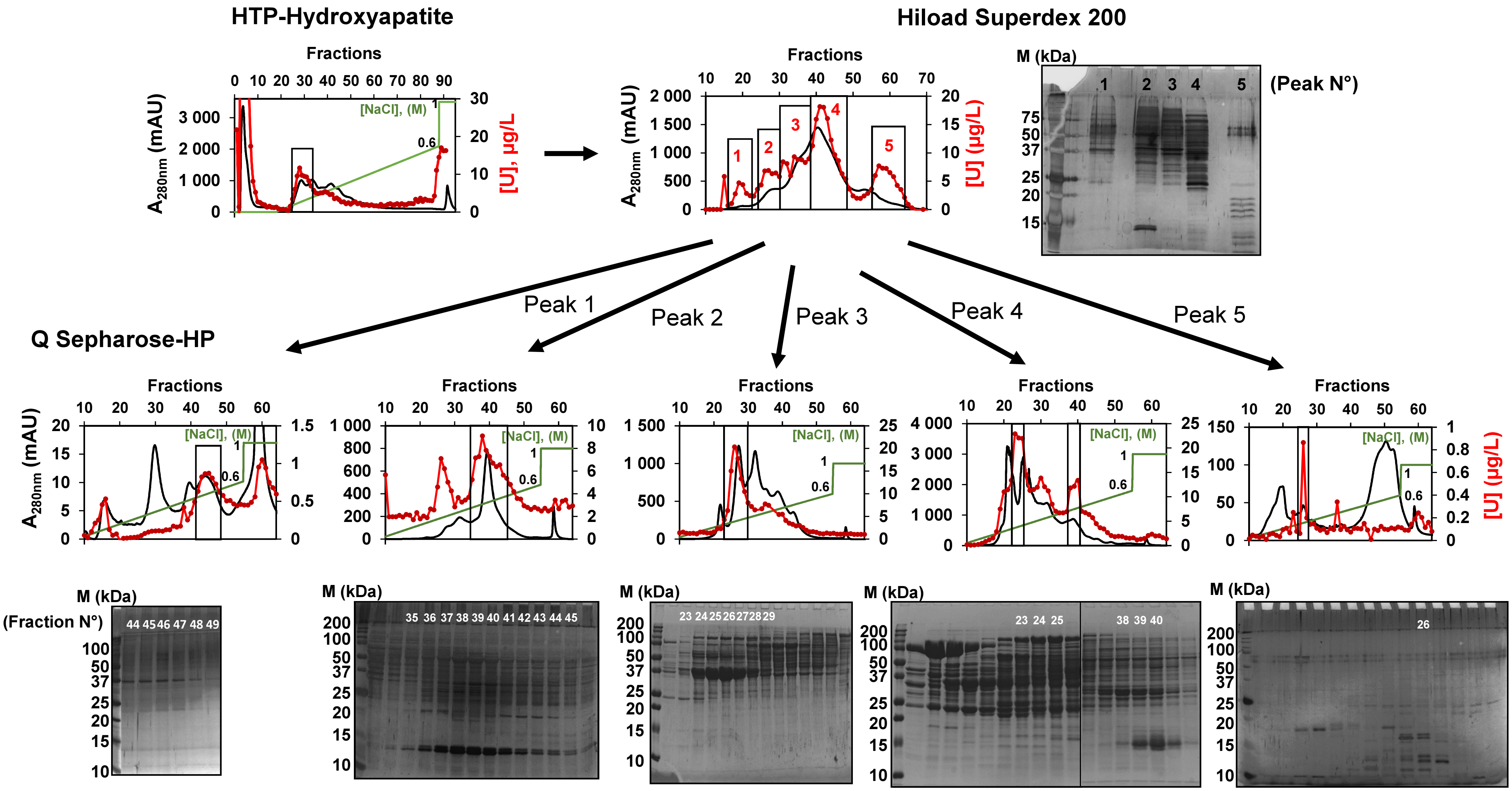


Supp. Figure S6 (see next page for legend)

**Supplementary Figure S6. Fractionation of UraBPs according to strategy 2.** Soluble proteins (250 mg in total) from 13 g of *A. thaliana* cells treated with 50 µM uranyl nitrate for 24 h, were separated on three successive chromatographic steps: an HTP hydroxyapatite column, a Hiload Superdex 200 column, and a Q-Sepharose HP column. Protein profiles are shown in black and U is shown in red. After the first hydroxyapatite step, a large U peak was recovered for fractionation on the Superdex 200 column. The U profile from this column was separated into five distinct peaks (1 to 5). Each of these peaks was loaded onto the Q-Sepharose column and the proteins were eluted with a 0 to 1 M NaCl salt gradient (shown in green). SDS-PAGE analyses of the eluted fractions (10 µL/well) are shown next to each profile. Fraction numbers of U peaks are given.

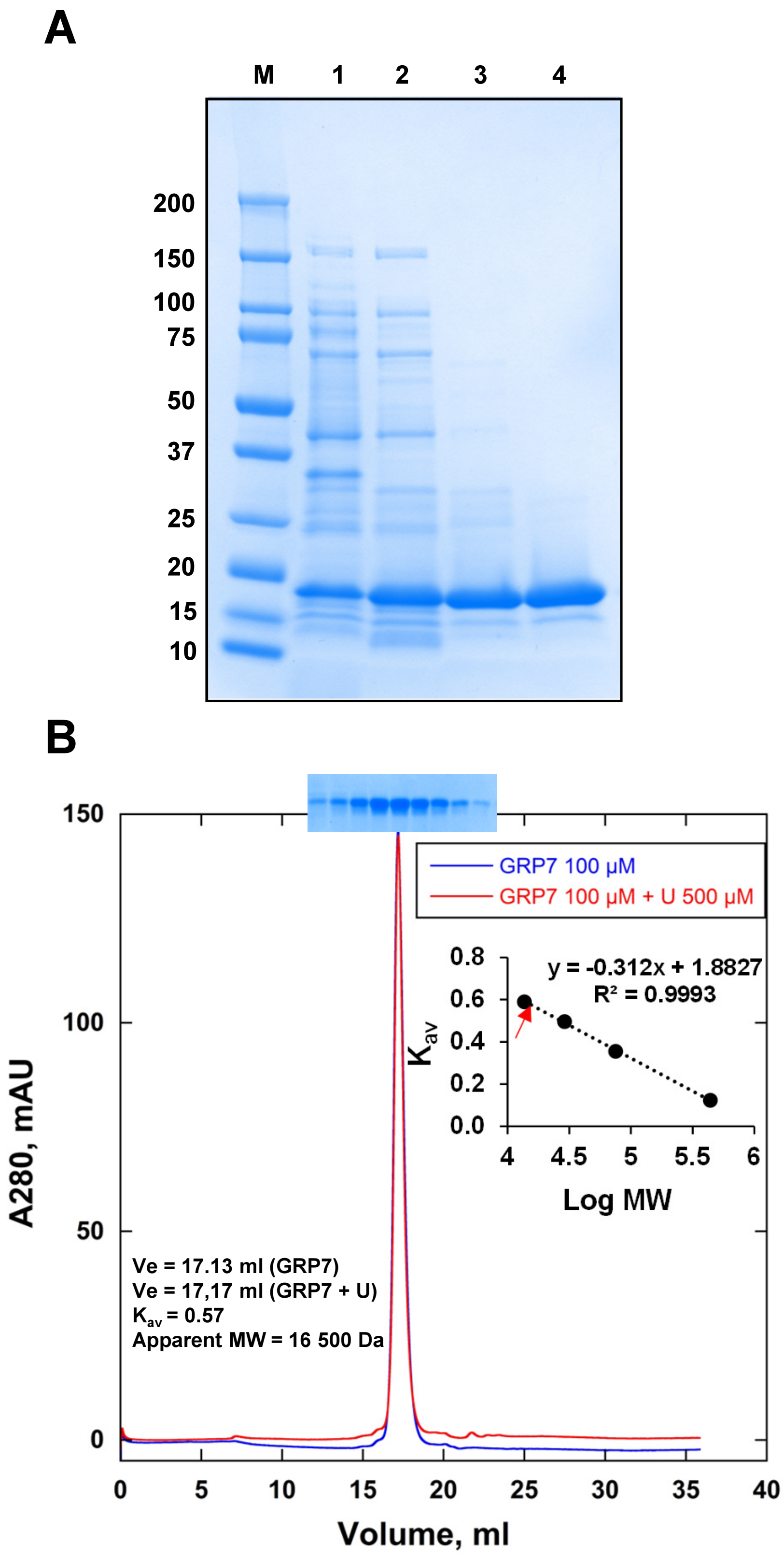


**Supp. Figure S7 (see next page for legend)**

**Supplementary Figure S7. Purification of recombinant GRP7 and determination of its oligomerization state.** **A**. Documentation of GRP7 purification. Polypeptides were separated by SDS-PAGE 12 % and stained with Coomassie Brilliant Blue. Lane 1: soluble proteins (25 µg) from *E. coli* Rosetta cells harboring pET28-GRP7 construct grown in the presence of IPTG; Lane 2: ammonium sulfate 40-60 % of saturation precipitating fraction (25 µg); Lane 3: Q-Sepharose column pool (10 µg); Lane 4: Superdex 75 column pool (8 µg); M, molecular mass markers. **B**. Apparent molecular mass estimation of native recombinant GRP7 and GRP7:U(VI) complex by gel filtration. Purified protein (100 µM) preincubated (in red) or not (in blue) with 500 µM uranyl nitrate was resolved by size exclusion chromatography onto a Superdex 200 Increase 10/300 GL column. Eluted fractions were analyzed by SDS-PAGE. Standard proteins for column calibration (inset) were ferritin (440 kDa), covalbumin (75 kDa), carbonic anhydrase (29 kDa), and ribonuclease A (13.7 kDa). *K*av = (Ve-Vo)/(Vt-Vo); Ve, elution volume; Vo, void volume; Vt, total volume.

**A**

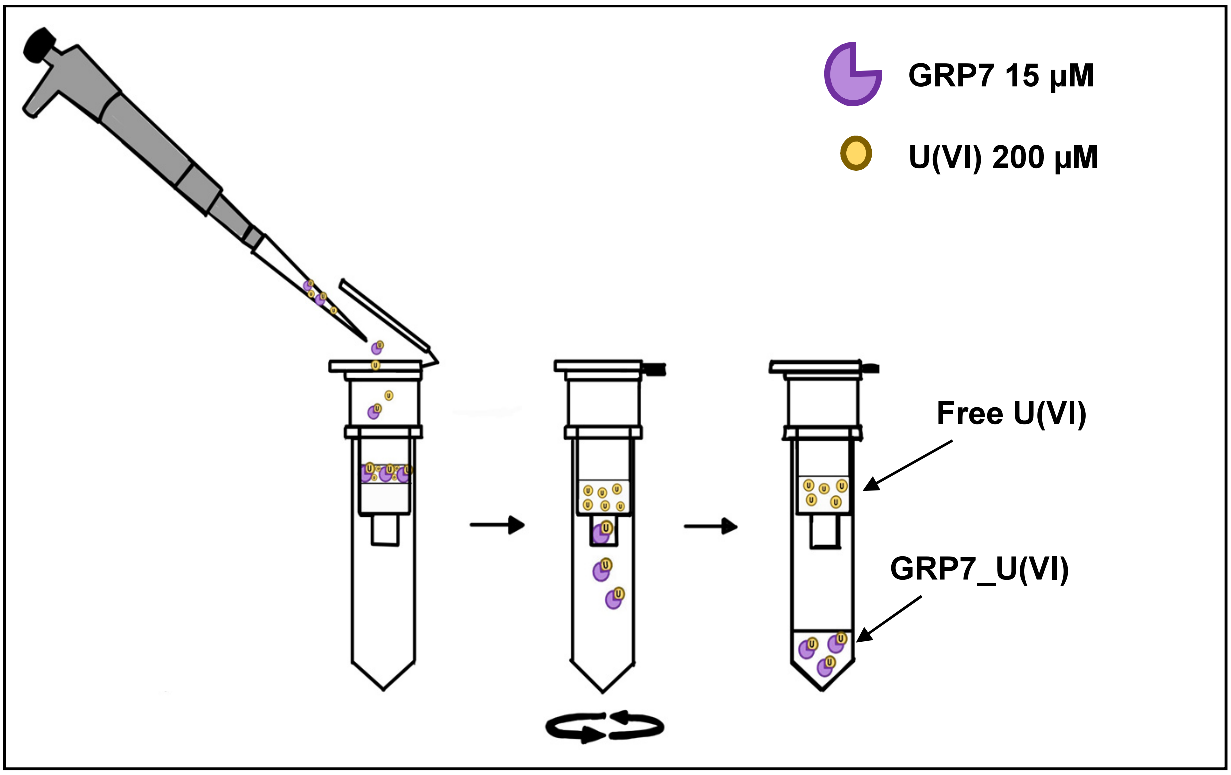


**B**

| Equivalent U in filtrate | | | | | | | |
|---|---|---|---|---|---|---|---|
| | *assay 1* | *assay 2* | *assay 3* | *assay 4* | | **Mean** | **SD** |
| **GRP7** | **2.47** | **2.19** | **1.97** | **1.94** | | **2.14** | **0.25** |
| **GRP7Δ** | **1.82** | **1.79** | **2.31** | **2.31** | | **2.05** | **0.27** |
| **GRP7ΔU1$_{mut}$** | **1.14** | **1.39** | **0.95** | **1.33** | | **1.20** | **0.18** |
| **GRP7ΔU2$_{mut}$** | **1.49** | **1.42** | **1.67** | **1.73** | | **1.57** | **0.13** |
| **GRP7ΔU1_U2$_{mut}$** | **0.31** | **0.23** | **0.22** | **0.24** | | **0.25** | **0.04** |

**Supplementary Figure S8. Determination of U(VI) binding to recombinant GRP7 protein variants. A.** Schematic illustration of the U(VI)-binding assay and the removal of unbound metal by centrifugal size exclusion chromatography. **B.** Quantification of U in filtrates (protein-U(VI) complexes) by the arsenazo III assay. Uranium to protein ratios are indicated. Detailed assay conditions are described in the Material and Methods section. Each assay was repeated four times.

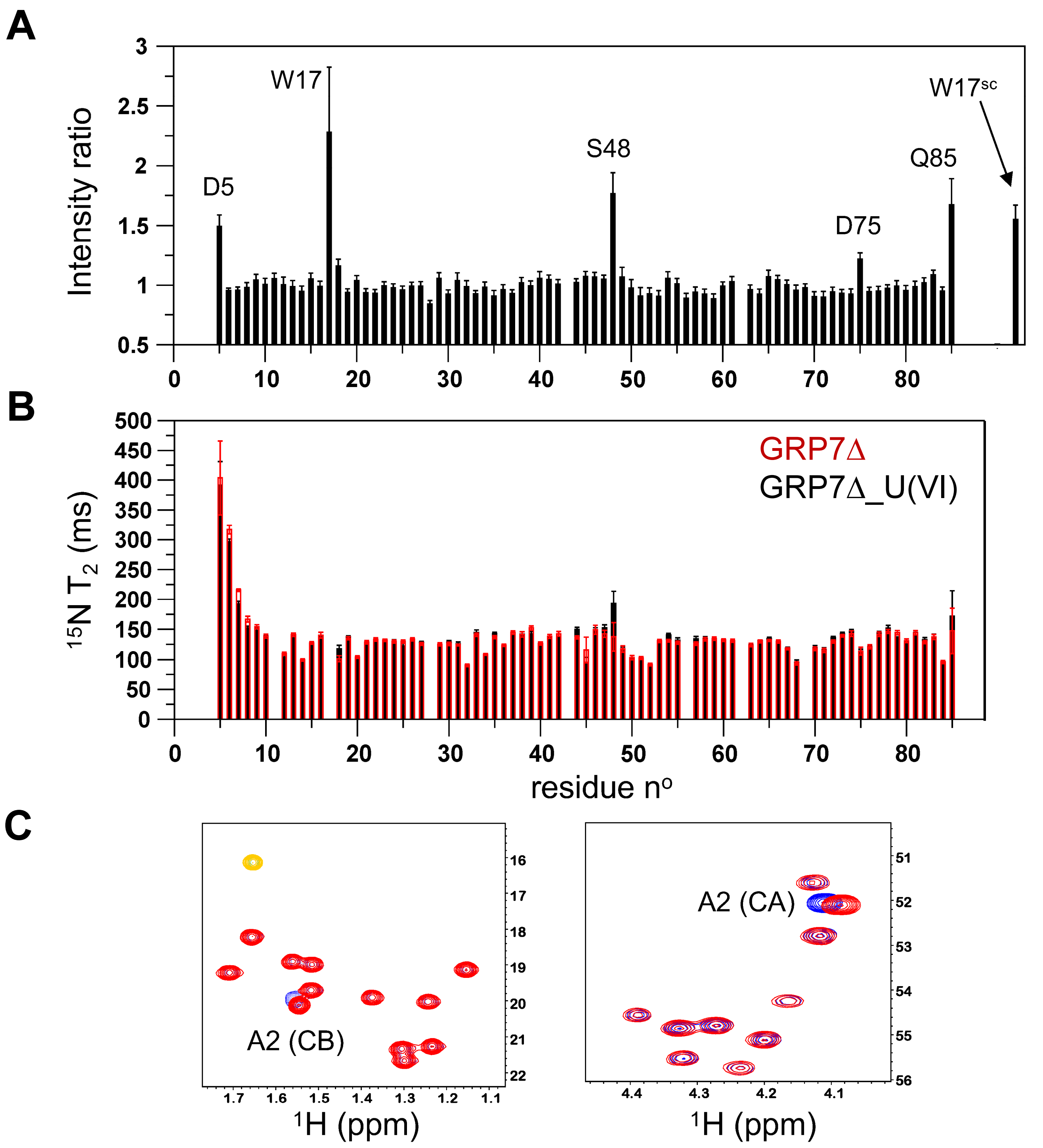


**Supplementary Figure S9. NMR characterization of the interaction between GRP7Δ and U(VI). A.** NMR peak intensity ratios (1:2 GRP7Δ:U complex / apo GRP7Δ) computed for individual amide sites and plotted as a function of protein sequence. **B.** $^{15}N$ relaxation time $T_2$ measured for the apo GRP7Δ (red bars) and a 1:2 GRP7Δ:U mixture (black bars). **C.** Superposition of $^{1}H$-$^{13}C$ spectral regions recorded for the apo GRP7Δ (red) and a 1:2 GRP7Δ:U mixture (blue). Similar to the full-length GRP7, the side chain resonances of Ala-2 show peak shifts upon uranyl binding.
